# Pla2g12b is Essential for Expansion of Nascent Lipoprotein Particles

**DOI:** 10.1101/2022.08.02.502564

**Authors:** James H. Thierer, Ombretta Foresti, Pradeep Kumar Yadav, Meredith H. Wilson, Tabea Moll, Meng-Chieh Shen, Elisabeth M. Busch-Nentwich, Margaret Morash, Karen L. Mohlke, John F. Rawls, Vivek Malhotra, M. Mahmood Hussain, Steven A. Farber

**Affiliations:** Department of Embryology, Carnegie Institution for Science, Baltimore, MD 21218, US; Department of Cell and Developmental Biology, Centre for Genomic Regulation, Barcelona 08003, ES; Department of Foundations of Medicine, NYU Long Island School of Medicine, Mineola, NY 11501, US; Department of Biology, Johns Hopkins University, Baltimore, MD 21218, US; School of Biological and Behavioural Sciences, Queen Mary University of London, London E1 4NS, GB; Department of Molecular Genetics and Microbiology, Duke University, Durham, NC 27708, US; Department of Genetics, University of North Carolina, Chapel Hill, NC 27599, US

**Keywords:** Lipoprotein, Apolipoprotein-B, Pla2g12b, Cardiovascular Disease

## Abstract

**SUMMARY:** Triglyceride-rich lipoproteins (TRLs) are micelle-like particles that enable efficient transport of lipids throughout the bloodstream, but also promote atherosclerotic cardiovascular disease. Despite this central relevance to cardiovascular disease, very little is known about how lipids are loaded onto nascent TRLs prior to secretion. Here we show that Pla2g12b, a gene with no previously described function, concentrates components of the TRL biogenesis machinery along the ER membrane to ensure efficient delivery of lipids to nascent TRLs. We find that the lipid-poor TRLs secreted in *PLA2G12B*^*-/-*^ mice and zebrafish support surprisingly normal growth and physiology while conferring profound resistance to atherosclerosis, and demonstrate that these same processes are conserved in human cells. Together these findings shed new light on the poorly understood process of TRL expansion, ascribe function to the previously uncharacterized gene Pla2g12b, and reveal a promising new strategy to remodel serum lipoproteins to prevent cardiovascular disease.

## INTRODUCTION

Triglyceride-Rich Lipoproteins (Feingold and Grunfeld, 2000) (TRLs) are composed of a core of hydrophobic lipids surrounded by an amphipathic coat of phospholipids and proteins(Davidson and Shelness, 2000) including the structural protein Apolipoprotein-B (ApoB) (Figure 1A). This structure enables TRLs to solubilize and transport large quantities of lipids through the aqueous circulatory system. TRLs produced by the liver are referred to as very-low density lipoproteins (VLDL), whereas those secreted from the intestine are called chylomicrons (Davidson and Shelness, 2000). Following secretion, TRLs are digested by intravascular lipases (Horton, 2019), resulting in a progressive reduction in lipoprotein size, density, and triglyceride content, and giving rise to the smaller lipoprotein species such as intermediate and low-density lipoproteins (IDL and LDL, Figure 1A). LDL particles can be cleared from the bloodstream via receptor-mediated endocytosis (Goldstein and Brown, 1974), but in some cases may penetrate the arterial wall and accumulate in the vascular endothelium. Ectopic lipoprotein accumulation is the initiating event of atherosclerotic cardiovascular disease (Tabas et al., 2007), which is one of the leading causes of morbidity and mortality worldwide (Lopez and Murray, 1998).

**Figure 1:**
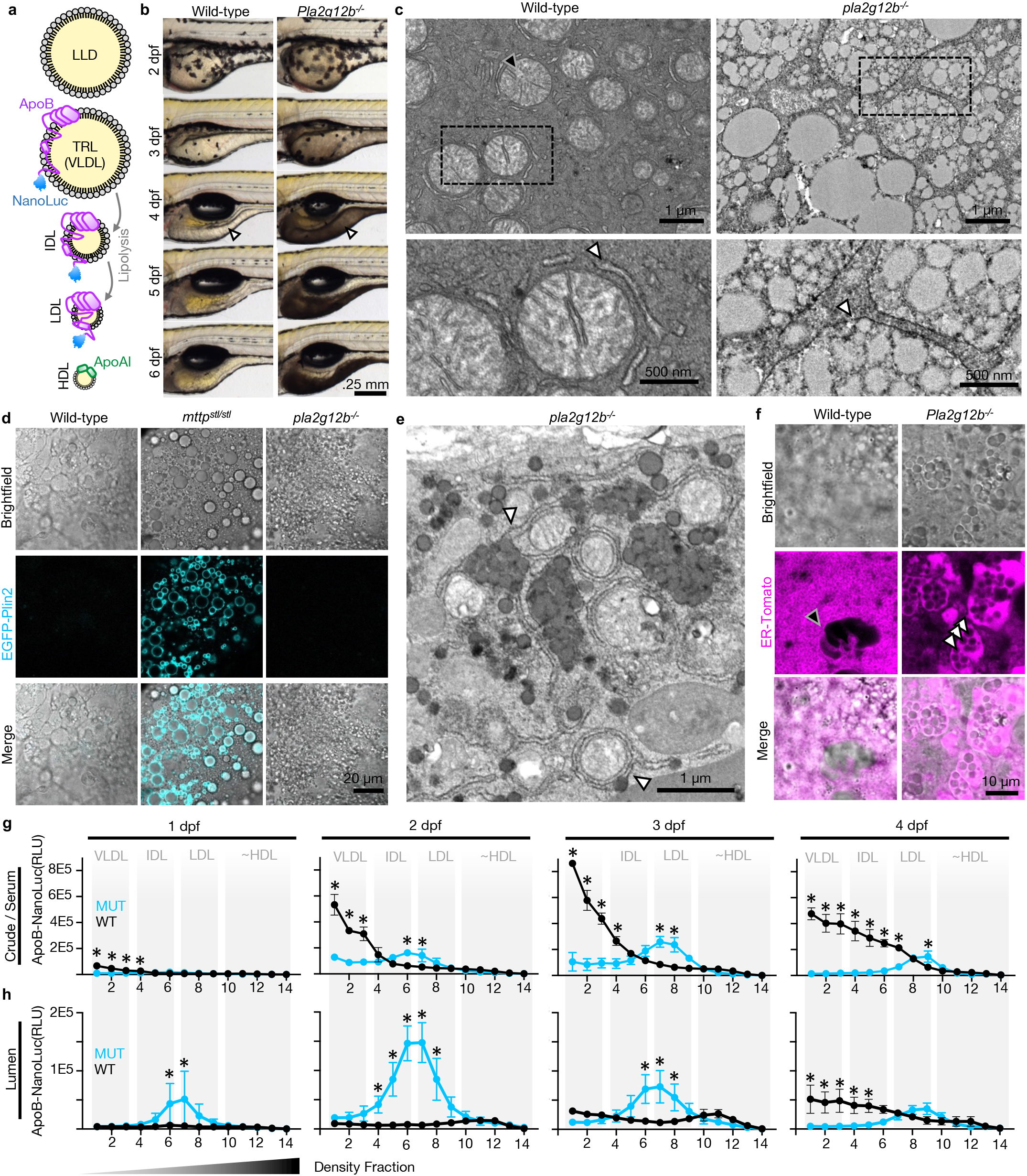
Lumenal lipid droplet accumulation and inefficient TRL lipidation in Pla2g12b mutants. **(a)** Schematic representation of lumenal lipid droplets (LLDs) which lack ApoB, various classes of ApoB-containing lipoproteins, and high-density lipoproteins (HDL) which contain ApoAI instead of ApoB. (**b)** Brightfield images of the embryonic yolk throughout larval zebrafish development (days post fertilization, dpf), showing delayed absorption and increased opacity (see arrowheads in 4 dpf panels) of *pla2g12b* mutant yolks. (**c)** Electron micrographs of the Yolk Syncytial Layer (YSL) at 4 dpf illustrating significant lipid accumulation in *pla2g12b*^*-/-*^ larvae. The YSL of wild-type larvae contains mitochondria (black arrowhead) surrounded by ER-tubules (white arrowhead in enlarged lower panel) with virtually no evidence of intracellular lipid accumulation. The YSL of *pla2g12b*^*-/-*^ larvae is dominated by lipid micelles that appear clustered together within a membranous organelle (white arrowhead in enlarged lower panel). (**d)** Confocal images of EGFP-Plin2 marker of cytoplasmic lipid droplets (CLDs) in wild-type, *mttp*^*stl/stl*^, and *pla2g12b*^*-/-*^ backgrounds showing that CLDs are virtually undetectable in *pla2g12b*^*-/-*^ larvae despite substantial intracellular lipid accumulation. (**e)** Electron micrograph of *pla2g12b*^*-/-*^ YSL at 3 dpf illustrating cases of ER-tubules expanding to accommodate clusters of lipid micelles (white arrowheads). (**f)** Confocal microscopy of the fluorescent ER-marker (tdTomato) confirms normal ER-morphology in wild-type controls (nucleus indicated by black arrowhead), while significant swelling of the ER lumen is apparent in *pla2g12b*^*-/-*^ larvae as a result of lipid inclusions (white arrowheads). (**g)** Density gradient profiles of the TRL marker ApoB-NanoLuc in crude larval zebrafish homogenate, which approximates the serum lipoprotein profile. Approximate densities of different lipoprotein classes are boxed in gray. Wild-type zebrafish (black line) produce buoyant VLDL particles early in development (1-2 dpf), which are lipolyzed into denser (LDL) particles by later stages of development (3-4 dpf). *pla2g12b* mutants show pronounced defects in the production of buoyant particles, and are enriched in denser (LDL-sized) particles at all stages of development (cyan line). (Two-way ANOVA p<.005 for interaction at all stages, asterisk denotes p<.05 by Šídák’s multiple comparison test). (**h)** Density gradient profiles of ApoB-NanoLuc in microsomal extracts throughout development, showing that *pla2g12b* mutants are unable to assemble buoyant TRLs within the ER lumen (Two-way ANOVA p<.005 for interaction at all stages, asterisk denotes p<.05 by Šídák’s multiple comparison test).

Despite their central role in cardiovascular disease, the pathways governing TRL biogenesis remain poorly understood. Lipoprotein biogenesis occurs within the ER lumen, where Microsomal Triglyceride Transfer Protein (Hussain et al., 2003) (Mtp) transfers lipids from the ER-membrane to ApoB to form a small nascent lipoprotein particle (Boren et al., 1994, Rustaeus et al., 1999, Demignot et al., 2014). However, the subsequent process of lipoprotein expansion and maturation remains relatively ill-defined. Mtp is capable of transferring additional lipids directly to the nascent TRL, but can also transfer lipids to ApoB-free lumenal lipid droplets (LLDs) (Wang et al., 2007). LLDs cannot be secreted on their own, but appear to be capable of fusing with nascent lipoproteins to generate lipid-rich TRLs (Cartwright and Higgins, 2001, Olofsson et al., 2000, Rustaeus et al., 1999). To date, it remains completely unknown how lipid flow and fusion events between LLDs and TRLs are regulated, and what cofactors govern these poorly understood processes.

Several lines of evidence suggest that Pla2g12b, a catalytically inactive member of the phospholipase A_2_ (PLA_2_) gene family (Rouault et al., 2003), may play a role in lipoprotein metabolism. Mutations in *pla2g12b* cause defects in lipoproteins homeostasis (Thierer et al., 2019, Guan et al., 2011) in both mice(Aljakna et al., 2012) and zebrafish (Kettleborough et al., 2013). The previously established zebrafish ENU allele *pla2g12b*^*sa659*^ disrupts an essential splice acceptor site in the last coding exon and induces nonsense-mediated decay (Anderson et al., 2017), and the mouse ENU allele *Pla2g12b*^*hlb218*^ causes a missense substitution in an essential cysteine residue (C129Y). For simplicity, mutant alleles will simply be referred to as *pla2g12b*^*-/-*^. Additionally, *Pla2g12b* expression is regulated by master regulators of lipid homeostasis in mice (Liu et al., 2016, Chen et al., 2019, Guan et al., 2011). Despite its many links to lipoprotein metabolism, the mechanistic role of Pla2g12b in lipoprotein metabolism has yet to be defined.

Using a combination of biochemical, genetic, and imaging techniques in a variety of model systems, we demonstrate that Pla2g12b serves to redirect ER-lipids away from un-secretable lumenal lipid droplets and towards secretable lipoproteins. Pla2g12b is thus an essential component of the lipoprotein expansion machinery, and the first known regulator of lipid partitioning between LLDs and TRLs within the ER-lumen.

## RESULTS

### Pla2g12b channels lipids to nascent lipoproteins

We analyzed the phylogenetic distribution of *pla2g12b* and found that it is conserved in every major vertebrate lineage, but absent in invertebrate lineages (Figure S1A). Our syntenic analyses suggest that *pla2g12b* emerged from a whole genome duplication event (Acharya and Ghosh, 2016) (Figure S1B), and its catalytically active ohnolog *pla2g12a* remains highly conserved as well (Figure S1A). We also find that *pla2g12b* expression is highly enriched in lipoprotein-producing tissues (Uhlen et al., 2015) (Figure S1C), and that expression is upregulated in response to food intake in the zebrafish intestine (Figure S1D). The phylogenetic, spatial, and temporal expression pattern of *pla2g12b* thus overlaps perfectly with TRL production, suggesting Pla2g12b may play an intracellular role in TRL biogenesis.

The larval zebrafish yolk-syncytial layer (YSL) presents a convenient system to monitor lipoprotein biogenesis, as this embryonic organ is highly specialized for TRL production, and is readily visualized via brightfield, fluorescent, and electron microscopy. We observed uncharacteristically dark or opaque YSLs in *pla2g12b*^*-/-*^ larvae (Figure 1B), which is indicative of intracellular lipid accumulation (steatosis) (Wilson et al., 2020). We then used electron microscopy to characterize the subcellular pattern of lipid accumulation within the YSL in more detail. Numerous lipid micelles were evident in electron micrographs in *pla2g12b*^*-/-*^ larvae, but were undetectable in wild-type controls (Figure 1C). Both the degree of yolk opacity and the number of lipid micelles increased throughout larval development (Figures S1E and S1F). Curiously, the pattern of lipid accumulation was not consistent with cytoplasmic lipid droplets (CLDs) that typically store excess cellular lipids (Sturley and Hussain, 2012, Farese and Walther, 2009, Guo et al., 2009). Instead, the majority of lipid micelles in *pla2g12b*^*-/-*^ larvae appeared to be bound within an endomembrane compartment (white arrowhead, Figure 1C).

Perilipins are cytosolic proteins that coat the surface of CLDs, and thus serve as highly specific molecular markers for CLDs (Wilson, 2021, Kimmel and Sztalryd, 2016). We used a fluorescent reporter Fus (*EGFP-plin2*) (Wilson, 2021) to confirm that CLDs are virtually undetectable in the YSL of *pla2g12b*^*-/-*^ larvae (Figure 1D). High-magnification electron micrographs indicated that even lipid droplets that appeared to be cytosolic in *pla2g12b*^*-/-*^ larvae were surrounded by an additional membrane (Figure S1G). These results suggest that in contrast to mutants with non-functional Mtp (*mttp*^*stl/stl*^) (Avraham-Davidi et al., 2012) which accumulate lipids in CLDs (Wilson et al., 2020) (Figure 1D), mutations in *pla2g12b* drive lipid accumulation almost exclusively within an endomembrane compartment.

To identify the endomembrane compartment containing the lipid micelles, we collected electron micrographs from an earlier time point when lipid accumulation was less pronounced, allowing other endomembrane structures to be visualized more easily (Figures 1E and S1F). At 3dpf, we identified clear examples of ER-tubules expanding to accommodate lipid micelles (white arrowheads, Figure 1E). Confocal microscopy was then performed on larvae carrying a fluorescent ER-marker (ER-TdTomato) to confirm localization of lipid micelles within ER-tubules. We observed extensive swelling and reorganization of the ER in *pla2g12b* mutants, as well as dark inclusions that overlap with lipid micelles visible in brightfield images (white arrowheads, Figure 1F), providing molecular evidence that lipid micelles accumulate within the ER lumen.

We then performed an epistasis experiment to determine whether Pla2g12b acts upstream or downstream of Mtp. If Pla2g12b acts upstream of Mtp, we would not expect inhibition of Mtp to have any effect on lipid accumulation in *pla2g12b* mutants. However, we found that the Mtp inhibitor Lomitapide was able to drive accumulation of CLDs and attenuate LLD accumulation in the *pla2g12b*^*-/-*^ background (Figure S1H), suggesting that Pla2g12b must act downstream of Mtp. We hypothesize that Mtp modulates lipid partitioning between the cytosolic and lumenal face of the ER membrane, and that Pla2g12b acts downstream of Mtp to regulate lipid flux within the ER lumen.

Two classes of lipid micelles are present within the ER lumen: Lumenal lipid droplets (LLDs) and nascent TRLs. These species can be differentiated by the protein marker Apolipoprotein-B (ApoB), which is present exclusively on the surface of TRLs (Figure 1A). We have previously established a chemiluminescent reporter of ApoB (ApoB-NanoLuc) called LipoGlo (Thierer et al., 2019), and reasoned that this reporter could be used in conjunction with cellular fractionation to monitor biogenesis of nascent TRLs in microsomal extracts.

We characterized the lipoprotein profile of both crude extracts (which are representative of serum lipoproteins) and microsomal contents from wild-type and *pla2g12b*^*-/-*^ larvae throughout the first 4 days of development (Figures 1G and 1H). The density cutoffs for different lipoprotein classes (Yee et al., 2008) are indicated with gray boxes. Note that particles with densities in the high-density lipoprotein range (∼HDL) are not *bona fide* HDL particles as they contain ApoB and not ApoAI (Figure 1A and 1H).

Consistent with previous findings, crude extracts from wild-type larvae were dominated by buoyant TRLs that are gradually lipolyzed into dense IDL/LDL particles (Figure 1G). By contrast, crude extracts from *pla2g12b*^*-/-*^ larvae contained significantly fewer lipoproteins overall, and exhibited depletion of buoyant particles and enrichment of dense species (Figure 1G). The lumenal lipoprotein profile of wild-type larvae contained two predominant species, a faint peak of HDL-sized lipoproteins that likely corresponds to newly formed TRLs that have not yet been lipidated (Boren et al., 1994, Rustaeus et al., 1999, Demignot et al., 2014), and a second peak of buoyant lipoproteins that represents more mature nascent VLDL particles (Figure 1H). In contrast, Pla2g12b mutants produce predominantly intermediate-sized particles with densities similar to LDL (Figure 1H).

The presence of unusually dense TRLs within both microsomal and crude serum extracts of *pla2g12b*^*-/-*^ larvae suggests that these mutants are fundamentally incapable of producing large TRLs. As the lipid micelles visible on electron micrographs reach diameters well above the expected diameters of VLDL particles (∼70 nm), we reason that they must be ApoB-free lumenal lipid droplets. Additionally, the number of lipid micelles visible in electron micrographs increases from 2-4 dpf (Figure S1F), whereas the yield of microsomal TRLs declines during this same period (Figure 1H). Thus, although we have yet to identify a specific molecular marker of LLDs, the opposing trends observed between ApoB-labeled TRLs and large lipid micelles visible on electron micrographs strongly suggests that the large lipid inclusions are ApoB-free LLDs.

### Pla2g12b is not essential for secretion of large cargoes

Whole-mount imaging of lipoprotein localization in larval zebrafish has conclusively demonstrated that lipoproteins are secreted into the serum of *pla2g12b*^*-/-*^ zebrafish (Thierer et al., 2019), but the buildup of nascent TRLs within the ER-lumen (Figure 1H) suggests that mild trafficking defects may exist in these mutants. We sought to investigate whether Pla2g12b plays a direct role in lipoprotein secretion, or whether the secretion defects may be a secondary consequence of abnormal lipid accumulation within the ER. To test this, we used the intracellular trafficking inhibitors Brefeldin-A (BFA) and Nocodazole (NOC) to disrupt TRL secretion in both wild-type and mutant larvae (Figure 2A-F). BFA interferes with anterograde ER-Golgi trafficking causing collapse of the Golgi apparatus (Figures 2A and 2B), and has previously been shown to disrupt TRL maturation in cultured cells by an unknown mechanism (Rustaeus et al., 1995). Exposure to BFA inhibited particle expansion in zebrafish larvae as well, and partially phenocopied the *pla2g12b*^*-/-*^ mutant phenotype (Figures 2C and 2D). However, BFA also led to a significant increase in the number of lumenal TRLs in both wild-type and mutant larvae (Figures 2C and 2D). The finding that BFA increases the yield of lumenal lipoproteins in both genotypes suggests that ER-to-Golgi trafficking of nascent lipoproteins is intact in *pla2g12b* mutants.

**Figure 2:**
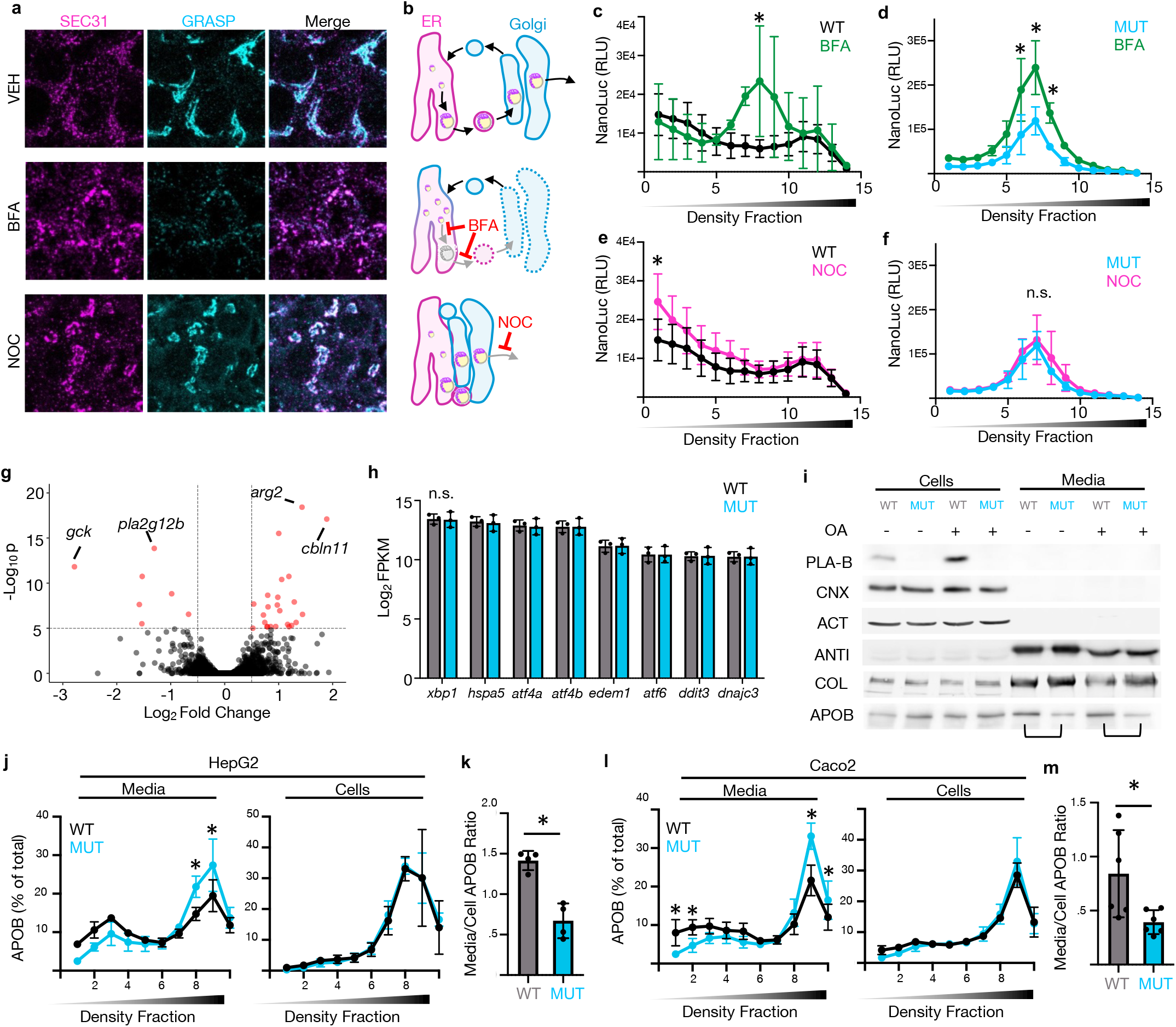
Lipoprotein secretion is intact in *pla2g12b* mutants. **(a)** Immunofluorescent images of co-localization between ER-exit sites (SEC31) and Golgi marker (GRASP65) in HepG2 cells in response to various trafficking inhibitors. Brefeldin-A (BFA) prevents anterograde ER-to-Golgi trafficking, causing dissolution of the Golgi and accumulation of Golgi-resident proteins in the ER. Nocodazole (NOC) depolymerizes microtubules, causing the Golgi to associate closely with the ER and inhibiting post-Golgi secretion. (**b)** Schematic illustration of the impact of trafficking inhibitors on ER (magenta) and Golgi (cyan) morphology. BFA causes the Golgi to collapse into the ER, while NOC blocks post-Golgi secretion and causes the Golgi to associate closely with the ER membrane. (**c)** Density profiles of ApoB-NanoLuc from microsomal extracts of wild-type zebrafish larvae exposed to the trafficking inhibitor Brefeldin-A (BFA), showing that this treatment partially phenocopies *pla2g12b* mutants by inhibiting both TRL expansion and secretion. (Two-way ANOVA p=.007 for effect of drug, asterisk denotes p<.05 by Šídák’s multiple comparison test). **(d)** BFA treatment does not affect the size distribution of nascent TRLs in mutant larvae, but does inhibit secretion (Two-way ANOVA p<.0001 for effect of drug, asterisk denotes p<.05 by Šídák’s multiple comparison test). (**e)** Nocodazole delays secretion of large TRLs from wild-type larvae. (Two-way ANOVA p=.0003 for effect of drug, asterisk denotes p<.05 by Šídák’s multiple comparison test). (**f)** NOC has no impact on the nascent TRL profile in mutant larvae. (Two-way ANOVA p=.19). **(g)** Volcano plot comparing transcriptomic sequencing of *pla2g12b* mutants to wild-type siblings. Thresholds set to p<1×10^−5^ and fold-change >1.41. (**h)** Genotype had no effect on expression of a panel of ER-stress markers. (Multiple t-tests, p>.7 for all pairwise comparisons). (**i)** Western blots monitoring impact of *pla2g12b* genotype on secretion of various proteins from HepG2 cells. PLA2G12B (PLA-B), CALNEXIN (CNX), and ACTIN (ACT) are not secreted. ANTI-TRYPSIN (ANTI) and COLLAGEN-XII (COL) are secreted normally irrespective of genotype, but APOB secretion appeared subjectively diminished in *PLA2G12B* mutants (black brackets). (**j)** Density profiles of APOB from the media of HepG2 cells normalized to total quantity of APOB secreted to highlight differences in density distribution. Mutant cells exhibit significant enrichment of dense TRLs in the media, but no significant changes are apparent in cell extracts. (Two-way ANOVA p<.0001 for interaction in media, p=.99 for interaction in cell extracts, asterisk denotes p<.05 by Šídák’s multiple comparison test). (**k)** Quantification of the media/intracellular APOB ratio shows significantly higher APOB secretion in WT cells. (Unpaired two-tailed t-test p=.0009). (**l)** Mutant Caco2 cells exhibit similar trends as HepG2 cells, with enrichment of dense TRLs and depletion of buoyant TRLs in the media. (Two-way ANOVA p<.0001 for interaction in media, p=.12 for interaction in cell extracts, asterisk denotes p<.05 by Šídák’s multiple comparison test) **(m)** Caco2 cells also exhibit reduced APOB secretion as measured by the ratio of APOB in cellular and media extracts. (Unpaired two-tailed t-test p=.025).

Nocodazole (NOC) depolymerizes microtubules, which disrupts post-Golgi vesicular trafficking and causes the Golgi to associate closely with the ER membrane (Figures 2A and 2B). Exposure to NOC significantly increased the yield of very-low density TRLs in wild-type, but not mutant, microsomal extracts (Figures 2E and 2F). We therefore consider NOC a useful probe to promote efficient capture of buoyant TRLs that are otherwise rapidly secreted from the ER and Golgi compartments. The absence of buoyant nascent TRLs in mutant microsomes exposed to NOC confirms that *pla2g12b* mutants cannot efficiently transfer lipids to nascent TRLs.

We hypothesized that if the secretory pathway were disrupted in *pla2g12b* mutants, it would lead to activation of an ER-stress response. We performed transcriptome-wide RNA-sequencing of *pla2g12b* mutants and their wild-type siblings at 4 days post-fertilization (Figure 2G, Table S1) and found no evidence of activation of ER-stress responsive genes (Vacaru et al., 2014) (Figure 2H).

To monitor the impact of Pla2g12b on TRL secretion more directly, CRISPR/Cas9 was used to generate stable knockouts in human liver (HepG2) and intestinal (Caco2) cell lines. Western blotting was then used to evaluate the secretion of various cargoes from wild-type and mutant cells (Figure 2I). PLA2G12B is undetectable in the media suggesting it is not secreted. Secretion of the small protein cargo Anti-Trypsin (ANTI) and the large protein cargo Collagen-XII (COL) was unaffected by *PLA2G12B* genotype, suggesting that both the canonical and large-cargo secretion pathways are intact in these mutants (Figure 2I). However, the media extracts of mutant cells contained significantly less APOB relative to wild-type samples (Figure 2I), which prompted us to investigate TRL secretion in more detail in these cell lines.

Consistent with prior results from zebrafish larvae, density-gradient ultracentrifugation revealed that buoyant TRLs were significantly depleted in the media extracts of mutant cell lines (Figures 2J and 2L). Note that these traces have been normalized to total APOB content to emphasize differences in density distribution. The proportion of APOB secreted into the media was significantly lower in *PLA2G12B* mutant cells (Figures 2K and 2M), confirming that although APOB secretion is intact, a mild defect is detectable in mutant cells. While there was a trend toward depletion of buoyant particles in cell extracts from both cell lines, this did not reach significance (Figures 2J and 2L), which we believe is attributable to low yield of buoyant TRLs from total cellular extracts.

The slightly reduced secretion of APOB observed in *PLA2G12B* mutants is therefore not the result of an inability to secrete TRLs, or disruptions in the canonical or large-cargo secretion pathways. We therefore propose that the trafficking aberrations observed are likely secondary effects resulting from abnormally small TRLs, an overabundance of large LLDs, or distortion of the ER.

### Binding Partners and Functional Domains of Pla2g12b

We next sought to characterize the mechanism by which Pla2g12b mediates TRL expansion, and reasoned that identifying its binding partners might provide clues regarding this mechanism. To investigate this, we generated a rescue plasmid that drives expression of bicistronic FLAG-tagged *pla2g12b* and GFP under control of a YSL-specific promoter (Wang et al., 2011) (Figure 3A). Microinjection of this vector into *pla2g12b* mutant larvae resulted in complete reversal of the darkened-yolk phenotype (Figure 3B), and GFP could be used as a fluorescent indicator to select successful injections. To identify potential binding partners, mutant zebrafish larvae were injected with rescue plasmids encoding FLAG-tagged *pla2g12b*, cross-linked in PFA, and subjected to unbiased co-immunoprecipitation and mass-spectrometry (Co-IP/MS) using anti-FLAG coated beads (Figure 3C).

**Figure 3:**
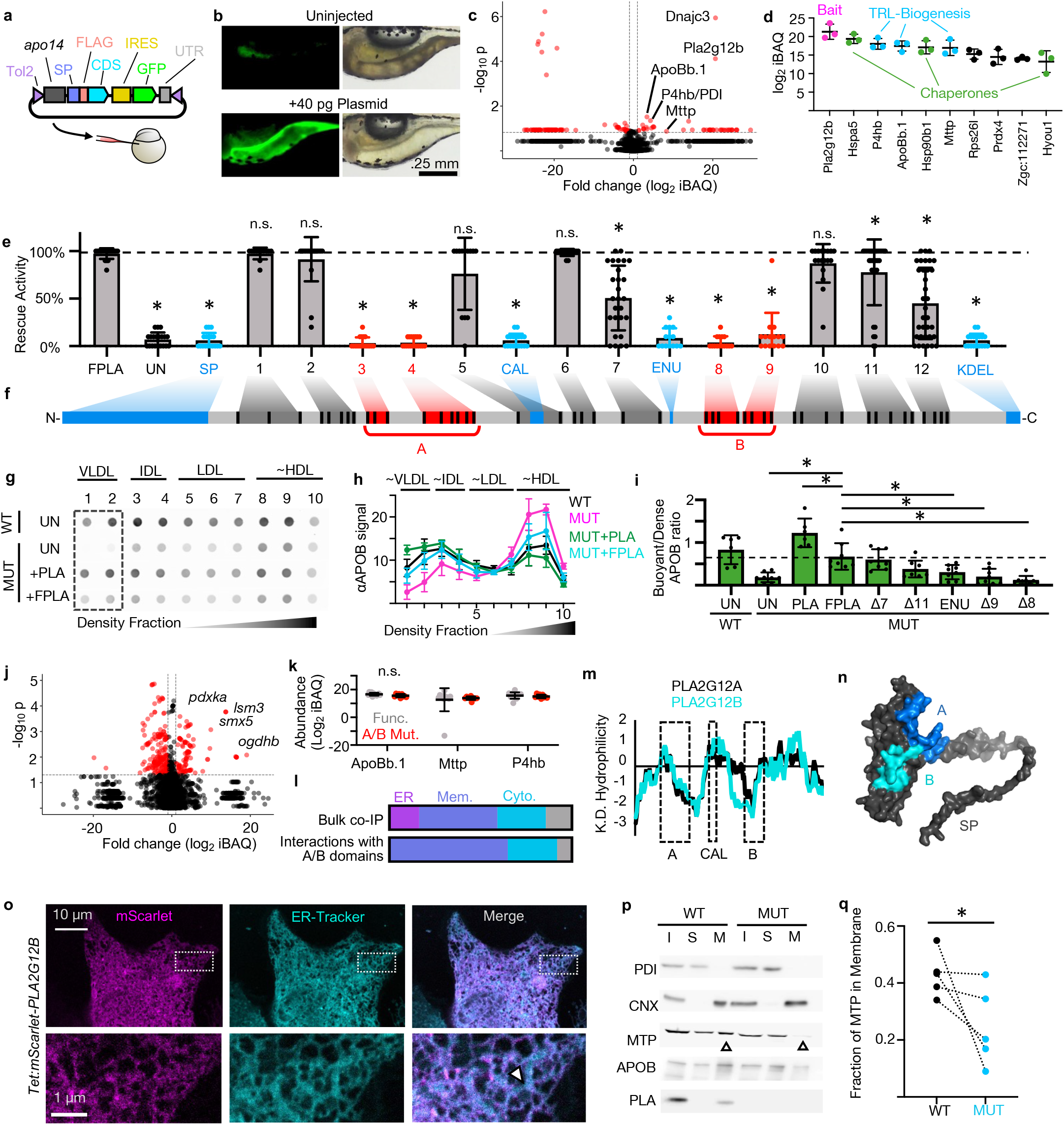
Membrane-interacting domains are essential to Pla2g12b function. **(a)** Schematic illustration of rescue plasmid encoding 3XFLAG-tagged *pla2g12b* coding sequence (CDS) and a GFP marker under control of a YSL-specific promoter (*apo14*). (**b)** Representative images of rescue assay, showing expression of the GFP marker and reversal of yolk-darkening in *pla2g12b* mutants injected with rescue plasmid. (**c)** Volcano plot of proteins detected in co-IP of FLAG-tagged Pla2g12b. Thresholds set to p<.15 and fold-change >2. (**d)** Most abundant proteins meeting significance and enrichment criteria in panel c, highlighting interactions between Pla2g12b (magenta) and core components of the TRL biogenesis pathway (light blue) and molecular chaperones (green). (**e)** Rescue activity of a panel of mutant alleles of *pla2g12b*. (One-way ANOVA p<.0001, asterisk denotes p<.05 by Tukey’s multiple comparison test). **(f)** Schematic of Pla2g12b primary structure highlighted to show amino acid substitutions (black bars) within in each allele. Domains predicted to have function *in silico (*blue) were all essential for function, as were several unannotated domains (red, labeled A and B). (**g)** Representative dot blot showing density distribution of APOB in media from cultured HepG2 cells showing that either endogenous or transgenic PLA2G12B is required to efficiently produce buoyant lipoproteins with densities similar to VLDL. (**h)** Quantification of APOB density distribution from dot blots. (**i)** Rescue activity of various Pla2g12b alleles quantified as buoyant/dense ratio, showing near wild-type rescue activity of 3xFLAG-tagged Pla2g12b (FPLA), and reduced rescue activity in alleles 8, 9, and ENU, which also compromised function in zebrafish rescue assays. (One-way ANOVA p<.0001, asterisk denotes p<.05 by Tukey’s multiple comparison test). **(j)** Volcano plot showing protein interactions unique to functional *pla2g12b* isoforms. Thresholds set to p<.05 and fold-change >4. (**k)** There was no significant difference in interactions with the core components of the TRL biogenesis machinery between functional and non-functional alleles of *pla2g12b*. (Multiple unpaired t-tests, p>.05 for all pairs). **(l)** Subcellular localization of top interacting partners in different co-IP/MS experiments, showing high enrichment of ER-resident proteins in the bulk pull-down, but surprising enrichment of membrane-associated and cytosolic proteins in pull-downs from functional Pla2g12b. (**m)** Hydropathy plot of Pla2g12b amino acid sequence (blue) overlaid with that of Pla2g12a (black), showing that the A and B domains are predominantly hydrophobic, whereas the calcium-binding domain is hydrophilic. (**n)** Mapping of functional domains onto predicted Pla2g12b structure. (**o)** Confocal images of colocalization between mScarlet-tagged PLA2G12B and ER-tracker dye in HepG2 cells highlighting localization of PLA2G12B to the periphery of the ER, while ER-tracker signal diffuses throughout the entire ER tubule or sheet (white arrowhead). (**p)** Western blots of various ER-resident proteins in input (I) soluble (S) and membrane (M) fractions collected from wild-type (WT) and *PLA2G12B* mutant HepG2 cell lines. PLA2G12B (PLA) and Calnexin (CNX) partition exclusively to the membrane fraction, and MTP is significantly enriched in the membrane fractions in wild-type extracts (White arrowheads). (**q)** Quantification of western blots shown in panel p, showing a enrichment of membrane-associated MTP in wild-type cell lines (Unpaired two-tailed t-test, p=.03).

Peptides were filtered for statistical significance and fold enrichment relative to pull-downs from uninjected (epitope-negative) larvae (Figure 3C, Table S2), and then ranked by abundance. The most abundant protein meeting these cutoff criteria was the bait protein Pla2g12b, validating epitope specificity in the pull-down (magenta, Figure 3D). The essential components of the TRL assembly pathway were among the most abundant proteins detected in the pull-down (cyan, Figure 3D), including the major zebrafish isoform of APOB (Otis et al., 2015) (ApoBb.1) and the two subunits of Mtp (Mttp and P4hb/PDI). We also detected a variety of chaperones (green, Figure 3D), and several miscellaneous proteins (black, Figure 3D). Based on these interactions, we can implicate Pla2g12b as a cofactor that participates directly in TRL biogenesis.

The functional domains of Pla2g12b have not been characterized. We reasoned that our rescue assay could be used to identify essential protein domains by introducing mutations into the *pla2g12b* coding sequence and evaluating their impact on rescue activity. We performed a literature review and *in silico* analyses to identify putatively functional protein domains (Figure S2A), and identified an amino-terminal signal peptide (SP), a putatively conserved calcium-binding motif (CAL), an essential Cysteine residue identified in an ENU mutagenesis screen in mice (ENU), and a carboxy-terminal ER-retention motif (KDEL) (blue labels, Figures 3E and 3F). We then used phylogenetic conservation to identify the amino acid positions most likely to have evolved new functional roles in Pla2g12b (Figures S2A and S2B). We selected residues that were both universally conserved in *pla2g12b* orthologs from various species (Zebrafish, Mouse, and Human), but had diverged from the closely related paralog *pla2g12a* (black bars, Figure 3F). We refer to this mutagenesis scheme as paralog recoding, which we believe maximizes the probability of identifying neofunctionalized residues while minimizing risk of inducing protein misfolding. As it was not practical to mutate each of these residues individually, we grouped them together to generate 12 alleles that contain blocks of adjacent substitutions (Alleles 1-12, Figures 3E and 3F).

Site-directed mutagenesis of the rescue plasmid was used to generate a panel of 16 mutant alleles: 4 alleles that disrupt protein domains predicted to have activity *in silico*, and 12 alleles generated through paralog recoding. Each allele was injected into *pla2g12b* mutant larvae and scored for its ability to reverse the dark-yolk phenotype (Figure 3E). Mutations in several domains were well-tolerated and had little or no impact on rescue activity (such as alleles 1, 2, 5, 6, and 10, gray labels, Figures 3E and 3F). By contrast, disruption of the domains predicted to have activity *in silico (*blue labels, Figures 3E and 3F) virtually abolished rescue activity. Four alleles generated by paralog recoding exhibited drastically reduced rescue activity (3, 4, 8, 9, red labels, Figures 3E and 3F). As these alleles occurred in adjacent pairs within the amino acid sequence, we thought it appropriate to group paired alleles together rather than discuss each as a distinct functional domain. Alleles 3 and 4 will be referred to as putative functional domain A, and alleles 8 and 9 corresponds to functional domain B (Figures 3E and 3F).

We then sought to evaluate whether our annotation of essential domains was conserved across species by repeating these rescue assays in cultured human cells. Mutant cell lines produced significantly fewer buoyant TRLs (Figures 2J and 2L), but transfection with plasmids that reintroduce *PLA2G12B* (+PLA) or FLAG-tagged Pla2g12b (+FPLA) are able to restore buoyant TRL production (Figures 3G and 3H). Rescue activity was quantified by calculating the ratio of APOB between the buoyant (1 and 2) and dense (8 and 9) fractions (Figures 3H and 3I). Consistent with results from the allelic series in zebrafish, alleles 7 and 8 showed comparable rescue activity to FLAG-tagged Pla2g12b, whereas alleles 8, 9, and ENU showed significantly reduced rescue activity.

We hypothesized that essential domains A and B may represent binding surfaces for ApoB and Mtp, and that Pla2g12b may promote lipid transfer to nascent TRLs by concentrating these proteins together into more stable complexes. To test this, we repeated co-IP/MS experiments using *pla2g12b* alleles with mutations in the A and B domains (Figure 3J, Table S3). Surprisingly, mutations in the A and B domains had no impact in interactions with ApoB or Mtp (Figure 3K). We thus performed an unbiased analysis of proteins that interact specifically with the A and B domains, and were surprised to find that these domains interact predominantly with membrane-associated and cytosolic proteins rather than those within the ER (Figure 3L).

The presence of an essential signal peptide and ER-retention signal indicate that Pla2g12b is likely localized within the ER, so we were surprised to see enrichment of membrane-associated and cytosolic proteins. This could be explained, however, if the A and B domains of Pla2g12b were associated with the ER-membrane. To investigate this possibility, we analyzed the hydropathy of Pla2g12b, and showed that the A and B domains are predominantly hydrophobic (Figure 3M). By contrast, the calcium-binding domain is situated in a hydrophilic stretch of the protein that likely faces the ER-lumen (Figure 3M). We then mapped the A and B domains onto the predicted structure of Pla2g12b generated by AlphaFold (Figures 3N and S3), and found that the A and B domains are clustered together on a hydrophobic face of the protein. These observations are consistent with the essential A and B domains mediating interactions with the hydrophobic lipids within the ER-membrane.

We then assessed the membrane association of Pla2g12b directly in human cultured cells. Confocal microscopy demonstrated significant colocalization of mScarlet-tagged PLA2G12B with ER-tracker dye, with PLA2G12B signal enriched in the periphery of the ER tubules (Figure 3O). Cultured cells were then fractionated into input (I), soluble (S) and membrane (M) fractions, and western blotting was used to evaluate the partitioning of various proteins between these fractions. Endogenous PLA2G12B was detectable exclusively in the membrane fraction (Figure 3P). Additionally, a significantly higher proportion of MTP was associated with the ER-membrane in wild-type cells (Figures 3P and 3Q).

The essential activities of Pla2g12b therefore include interactions with APOB, MTP, molecular chaperones, and the ER-membrane. We also annotate five essential features within the Pla2g12b protein, including a signal peptide, an ER-retention signal, a calcium binding domain, and two hydrophobic domains that mediate interaction with the ER membrane. These findings suggest that Pla2g12b promotes TRL expansion by concentrating proteins required for lipoprotein biogenesis along the ER-membrane, which is also the site of neutral lipid synthesis.

### Calcium promotes TRL expansion *in vitro* and *in vivo*

As the calcium-binding domain of Pla2g12b is essential for function, we predicted that depletion of ER-calcium with Thapsigargin (THAP) would also inhibit TRL expansion. As expected, THAP treatment led to accumulation of poorly lipidated nascent TRLs in wild-type embryos, representing a partial phenocopy of Pla2g12b mutants (Figure 4A). As discussed earlier, NOC can be used to promote efficient capture of buoyant TRLs in microsomal extracts (Figure 4B). However, co-treatment with THAP blocked the accumulation of buoyant lipoproteins driven by NOC exposure, and instead resulted in an increased yield of abnormally dense nascent TRLs (Figure 4C). These findings support an essential role of calcium in TRL expansion *in vivo*.

**Figure 4:**
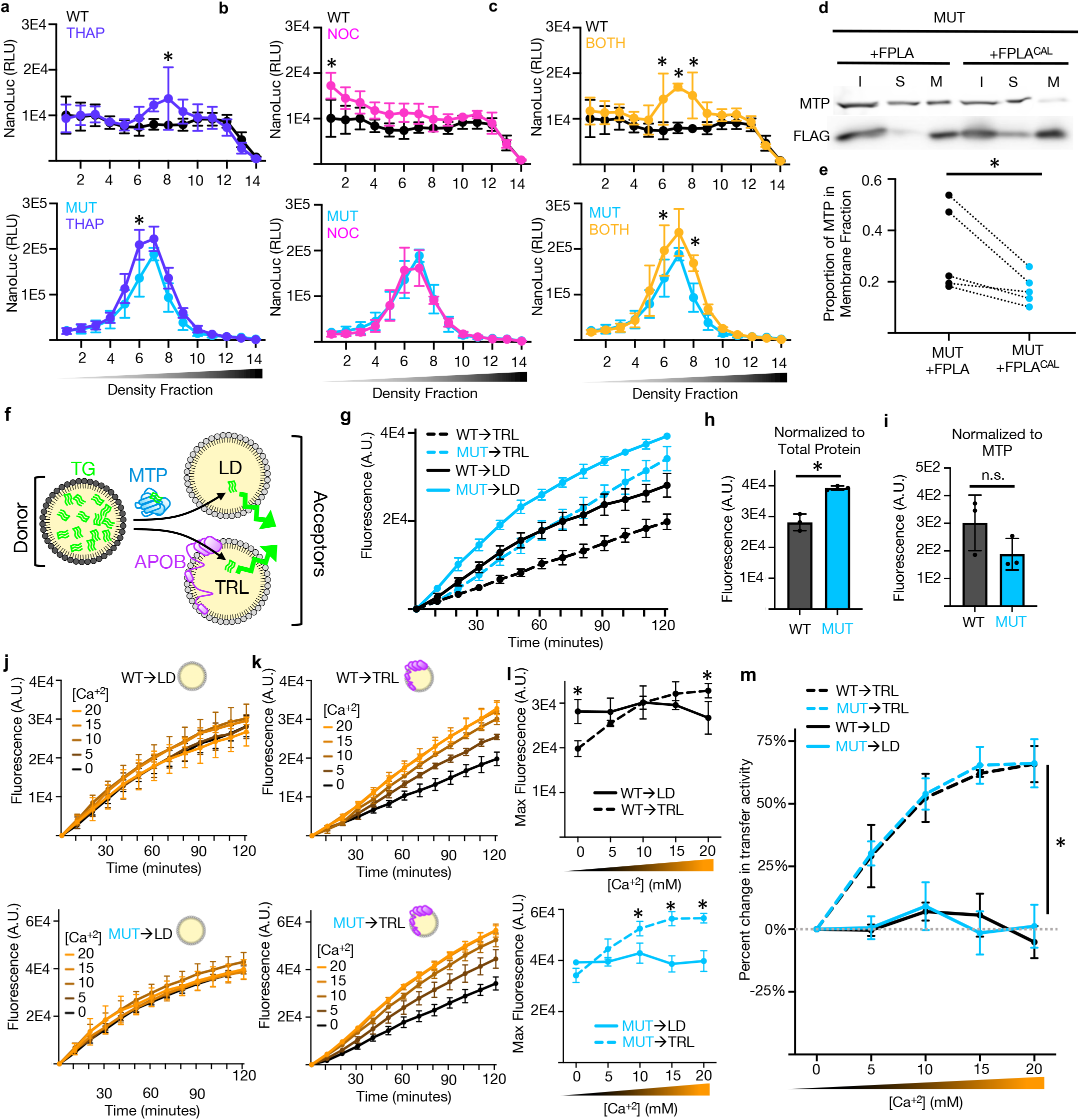
Calcium enables TRL expansion *in vivo* and *in vitro*. **(a)** Density distribution of lumenal ApoB in wild-type (upper panel) and mutant (lower panel) larvae exposed to thapsigargin (THAP) to deplete ER-calcium. Calcium depletion with THAP drove accumulation of unusually dense nascent lipoproteins in wild-type larvae. (Two-way ANOVA p<.005 for interaction for both genotypes, asterisk denotes p<.05 by Šídák’s multiple comparison test) **(b)** Nocodazole (NOC) exposure increases the yield of buoyant lipoproteins in lumenal extracts of wild-type, but not mutant, larvae. (Two-way ANOVA p<.005 for interaction in wild-type extracts, asterisk denotes p<.05 by Šídák’s multiple comparison test) **(c)** Co-treatment with both Thapsigargin and Nocodazole increases the accumulation of dense TRLs in microsomal extracts confirming the importance of calcium in TRL expansion. (Two-way ANOVA p<.005 for interaction for both genotypes, asterisk denotes p<.05 by Šídák’s multiple comparison test) **(d)** Representative western blot of input (I), soluble (S), and membrane (M) extracts from HepG2 cells demonstrating that partitioning of MTP and PLA2G12B to the membrane fraction is greatly reduced following transfection with a calcium-binding mutant of *PLA2G12B (*+FPLA^CAL^) when compared to wild-type FLAG-tagged *PLA2G12B* (+FPLA). (**e)** Quantification of western blots described in panel d, showing significantly reduced membrane association of MTP in the presence of the calcium-binding mutant of *PLA2G12B (*+FPLA^CAL^). (Two-tailed ratio-paired t-test, p=.008). (**f)** Schematic representation of *in vitro* triglyceride transfer assay, showing how MTP transfers fluorescently labeled triglycerides from a quenched environment (donor vesicle) to an unquenched acceptor vesicle or VLDL particle to generate a fluorescent signal. (**g)** Triglyceride transfer rates were higher in microsomal extracts from *PLA2G12B* mutant mice (**h)** when total protein was normalized between extracts, but (**i)** this difference disappeared when extracts were normalized to MTP content. (**j)** Triglyceride transfer to APOB-free vesicles was unaffected by calcium supplementation in either wild-type (upper panel) or mutant (lower panel) extracts (Two-way ANOVA p>.5 for both genotypes). (**k)** Calcium supplementation significantly increased transfer activity in both wild-type (upper panel) and mutant extracts (lower panel) when VLDL particles were used as acceptors. (Two-way ANOVA p<.001 for both genotypes). (**l)** Comparison of total transfer activity in wild-type (upper panel) and mutant (lower panel) extracts at various concentrations showing that transfer to APOB-free vesicles is favored under low-calcium conditions, but shifts to favor TRL acceptors in the presence of hyperphysiological calcium. (Two-way ANOVA p<.001 for both genotypes, asterisk denotes p<.05 by Šídák’s multiple comparison test). (**m)** Plot of percent change in transfer activity in response to calcium supplementation for for different genotypes and acceptor vesicles, showing that calcium increases transfer activity to VLDL acceptors by ∼65% but has no impact on transfer to APOB-free micelles. (Two-way ANOVA p<.001 for both genotypes, p<.0001 by Šídák’s multiple comparison test for LD vs. TRL acceptors at all time points, no significant differences between genotypes).

We next evaluated whether disruption of the calcium-binding segment of PLA2G12B altered the sub-organellar localization of MTP. Western Blotting revealed that a significantly lower proportion of MTP partitioned to the membrane fraction when the calcium-binding segment of *PLA2G12B* was disrupted (Figure 4D and 4E), suggesting that this activity may be calcium-dependent.

We then used an established *in vitro* assay (Athar et al., 2004) to directly measure the impact of calcium and *PLA2G12B* genotype on triglyceride transfer rates. Triglyceride transfer was monitored by quantifying the increase in fluorescence as MTP transfers fluorescently labeled triglycerides from quenched donor particles to unquenched acceptor particles (Figure 4F). The standard *in vitro* transfer assay uses APOB-free lipid micelles as acceptors, but APOB-containing lipoproteins also function as acceptor particles that better recapitulate the TRL expansion process (Athar et al., 2004) (Figure 4F).

Microsomal extracts were collected from wild-type and *PLA2G12B*^*-/-*^ mouse livers and used as a source of MTP. We were initially surprised to see higher triglyceride transfer activity in mutant extracts (cyan, Figure 4G and 4H) for both APOB-free and TRL acceptor particles, but found that this effect disappeared when extracts were normalized to MTP content rather than total protein content (Figure 4I). Further investigation will be required to discern whether disruption of *PLA2G12B* leads to hepatic overexpression of MTP, or simply increases MTP yield by favoring partitioning to the soluble fraction as we observed in cultured cells (Figure 4D).

Triglyceride transfer activity was then measured in the context of varying concentrations of calcium, ranging from 0 to 20 mM. Calcium supplementation had no impact on transfer to APOB-free vesicles (Figure 4J), but showed dose-dependent enhancement of transfer to VLDL acceptors (Figure 4K). Triglyceride transfer rates are higher for APOB-free acceptor particles in low calcium conditions, but in the context of 20mM calcium this trend reverses and transfer to VLDL particles becomes more efficient (Figure 4L). Calcium increased transfer activity to VLDL acceptors ∼65% irrespective of *PLA2G12B* genotype (Figure 4M). However, as PLA2G12B partitions to the insoluble membrane fraction, we would expect it to pellet with the membrane fraction of microsomal extracts and thus be unavailable to participate in the aqueous *in vitro* transfer reaction described here.

### Pla2g12b mutants exhibit normal growth and physiology

We next sought to characterize the broader physiological impacts of disrupted Pla2g12b function. We observed significantly slower growth in *pla2g12b* mutant zebrafish in the larval stages (Figure 5A), which likely results from significant retention of yolk nutrients in the YSL. Conversely, we were unable to detect any difference in body size in adult zebrafish (Figure 5B), suggesting that this growth defect is transient and is most pronounced in the lecithotrophic stages. Although homozygous *pla2g12b* mutants were slightly underrepresented in progeny from three separate incrosses of heterozygous parents, this distribution did not differ significantly from expected mendelian ratios (p=.06, Figure 5C) suggesting that there is not an appreciable defect in survival. Lipid accumulation has previously been reported in lipoprotein-producing tissues of *Pla2g12b* mutant mice, and we observed similar patterns of lipid accumulation in both the liver (Figure 5D) and intestine (Figure 5E) of adult zebrafish.

**Figure 5:**
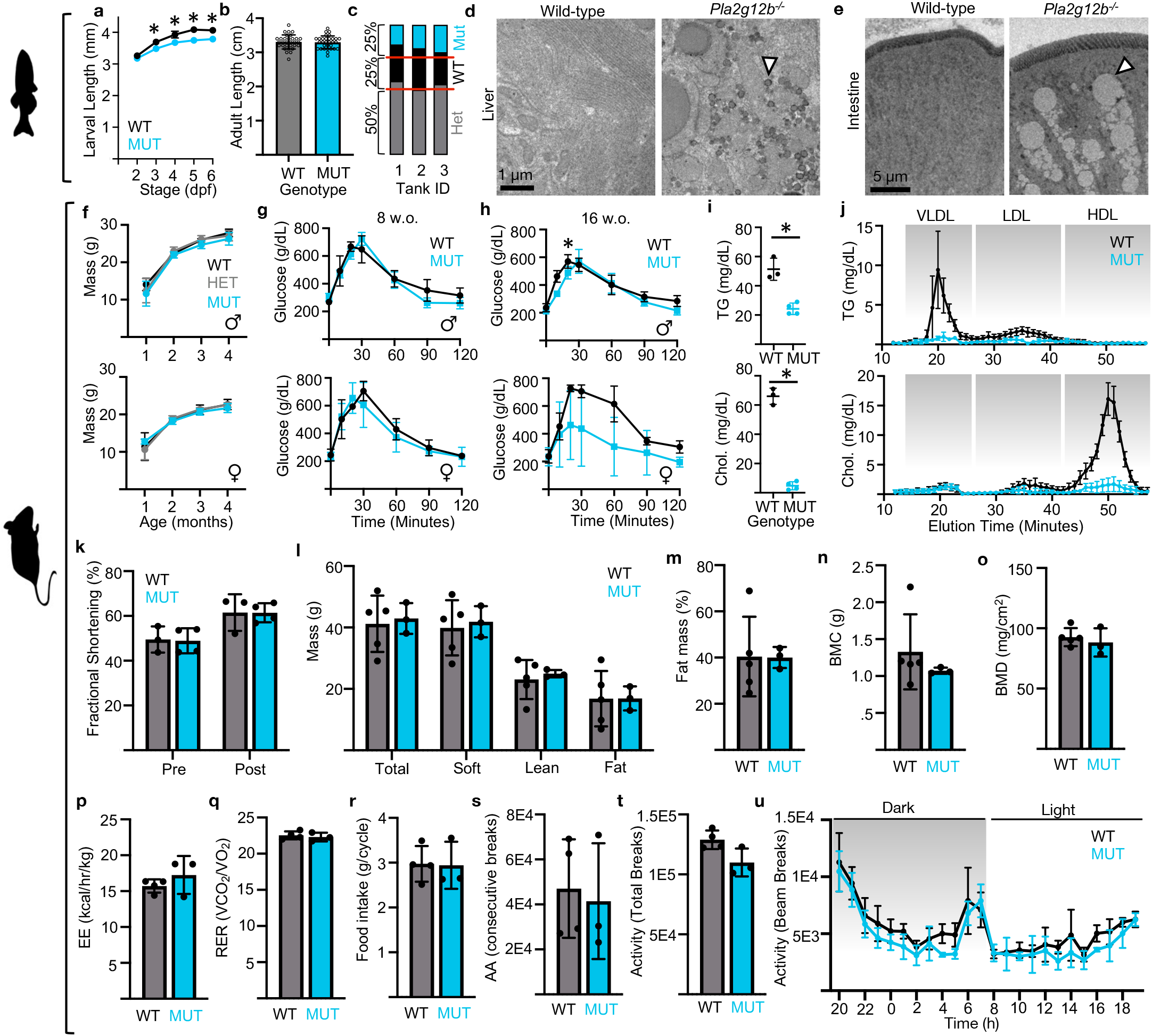
Mutations in Pla2g12b cause mild disruptions to systemic metabolism. **(a)** Body length is slightly reduced in *pla2g12b* mutant larvae. (Two-way ANOVA p<.0001 for interaction, asterisk denotes p<.0001 by Šídák’s multiple comparison test). (**b)** Adult body size was not significantly different between *pla2g12b* mutant zebrafish and wild-type siblings. (Unpaired two-tailed t-test, p=.80). (**c)** There was a trend towards depletion of homozygous *pla2g12b* mutants surviving to adulthood when raised with sibling controls, but ratios were not significantly different from expected mendelian ratios across three independent clutches. (Unpaired two-tailed t-test, p=.061). (**d)** Electron micrographs of liver and (**e)** intestinal sections from adult zebrafish are indicative of excess lipid accumulation in lipoprotein-producing tissues. (**f)** *PLA2G12B* genotype had no effect on body weight in male (upper panel) or female (lower panel) mice. (Two-way ANOVA p>.5 for both sexes). (**g)** Glucose tolerance was not significantly different in 8 week-old mice. (Two-way ANOVA p>.2 for both sexes). (**h)** Differences in glucose tolerance were detected in both sexes by 16 weeks of age. (Two-way ANOVA p<.005 for both sexes, asterisk denotes p<.05 by Šídák’s multiple comparison). (**i)** Serum triglycerides (upper panel) and cholesterol (lower panel) were significantly reduced in *PLA2G12B* mutants. (Unpaired two-tailed t-test, p<.005). (**j)** FPLC traces demonstrate that the lipid content of all lipoprotein classes, including HDL, are reduced in *PLA2G12B* mutants. (Two-way ANOVA p<.0001). (**k)** Quantification of fractional shortening in mice subjected to a dobutamine stress test, showing there is no defect in cardiac function in *pla2g12b* mutants. (One-way ANOVA p>.9 for effect of genotype). (**l)** Body composition parameters from DEXA scan show no differences in total, soft, lean, or fat mass, nor (**m)** fat mass percentage, (**n)** bone mineral content, or (**o)** bone-mineral density. (Unpaired two-tailed t-test, p>.05) (**p)** Quantification of parameters from CLAMS assay showed no difference in energy expenditure, (**q)** respiratory exchange rate, (**r)** food intake, or (**s)** ambulatory activity. (**t)** A slight reduction in total activity was detected in mutant males, but (**u)** was not restricted to a particular time of day. (Panels p-t: Not significant by unpaired t-test, p>.05. Panel u: Unpaired t-test p=.047, not significant after adjustment for multiple comparisons. Genotype had a significant effect on total activity by Two-way ANOVA, p<.0001, *post hoc* Šídák’s test used for multiple comparisons was not significant at any individual time point).

We also performed a broad panel of physiological and metabolic profiling in *PLA2G12B* mutant mice. We did not detect a growth defect in males or females (Figure 5F). We suspected that hepatic steatosis and serum hypolipidemia might result in broad changes in systemic nutrient metabolism, but surprisingly found no difference in glucose tolerance in 8-week-old mice (Figure 5G). We repeated experiments at 16 weeks and detected increased glucose tolerance in *PLA2G12B* mutants, suggesting that *pla2g12b* genotype may interact with age to influence systemic glucose metabolism (Figure 5H). We also performed extensive lipoprotein profiling of *PLA2G12B* mutants, and detected profound reductions in serum cholesterol and triglycerides (Figure 5I) that were distributed across all lipoprotein classes (Figure 5J).

TRLs are an important source of fuel for cardiomyocytes (Yu et al., 2005). Echocardiography was used to assess heart function in the context of a dobutamine stress-test, but *PLA2G12B* genotype had no impact on fractional shortening (Figure 5K) or any of the other cardiac parameters tested. Dual Energy X-Ray Absorptiometry (DEXA) scans were used to assess body composition, which was also unaffected by *PLA2G12B* genotype (Figures 5L-5O). A Comprehensive Lab Animal Monitoring System (CLAMS) was then used to assess numerous metabolic and activity parameters over a 24-hour window. None of these parameters were significantly affected by *PLA2G12B* genotype (Figures 5P-5U) after adjusting significance thresholds to account for multiple tests.

### Disruption of Pla2g12b confers resistance to atherosclerosis

We hypothesized that the near total absence of serum lipoproteins in *PLA2G12B* mutants would provide resistance to atherosclerotic cardiovascular disease. To test this, mice were subjected to a pro-atherosclerotic disease paradigm (Kumar et al., 2017) involving Adeno-associated virus (AAV)-mediated overexpression of Proprotein Convertase Subtilisin/Kexin Type 9 (PCSK9) and high-fat diet for a period of 20 weeks. PCSK9 is a circulating protein that promotes turnover and degradation of the LDL-receptor (Cohen et al., 2005), so overexpression of this protein drives substantial increases in serum lipoprotein levels by interfering with receptor-mediated lipoprotein clearance. Body weight increased significantly over the trial period, but remained indistinguishable between genotypes (Figure 6A). Serum triglycerides and cholesterol levels increased dramatically in both genotypes, but remained significantly higher in wild-type mice (Figure 6B and 6C). FPLC was used to fractionate serum lipoproteins at the end of the trial period, and demonstrated higher lipid content in wild-type animals across all lipoprotein classes (Figure 6D and 6E).

**Figure 6:**
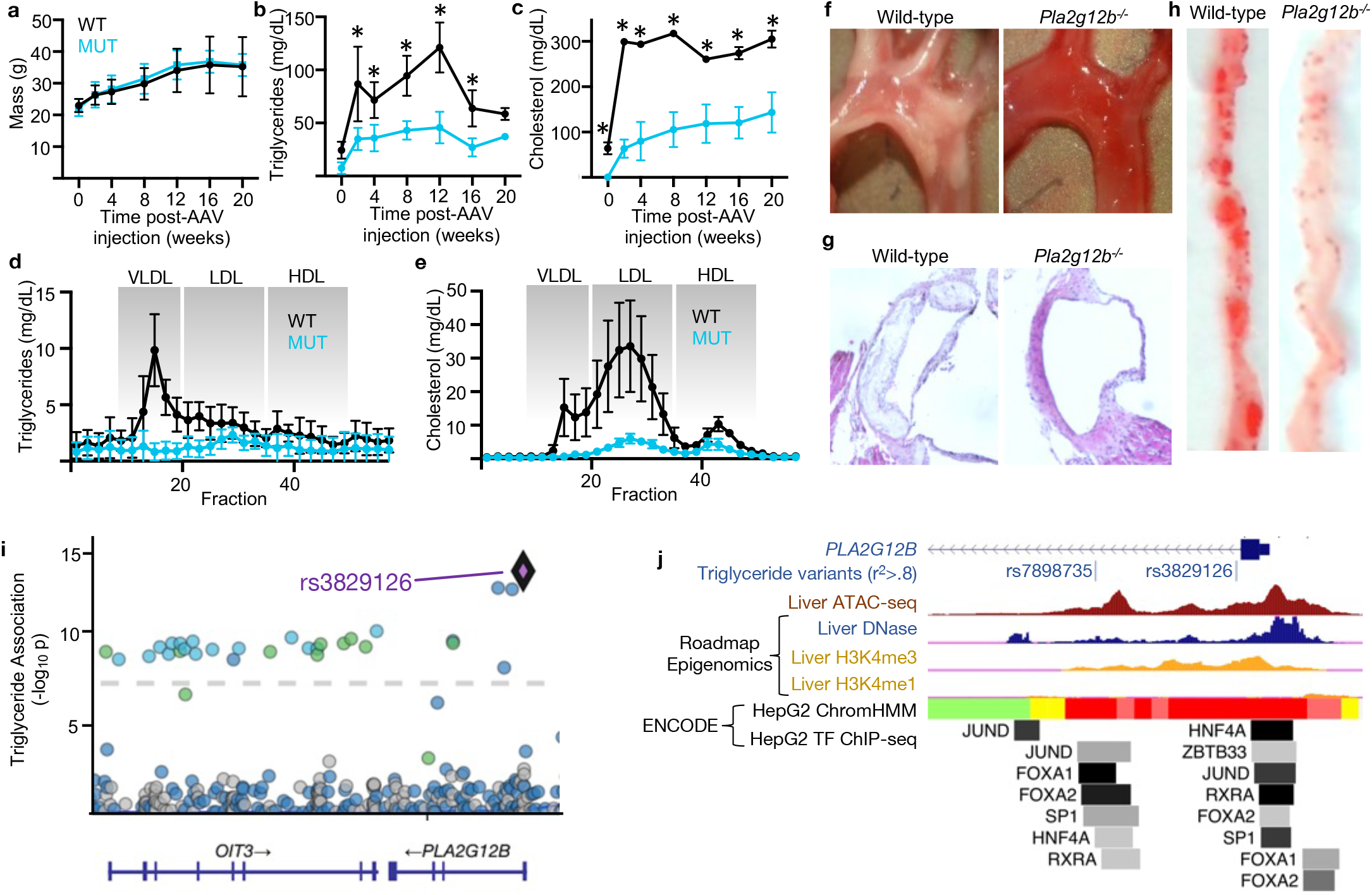
Pla2g12b mutants are resistant to hyperlipidemia and atherosclerosis. **(a)** Initiation of a pro-atherosclerotic paradigm did not induce divergence of body weights between *PLA2G12B* mutants and wild-type siblings. (Two-way ANOVA, p=.63). (**b)** Plasma triglycerides and (**c)** cholesterol levels increased significantly in both genotypes in response to the pro-atherosclerotic paradigm, but serum lipid levels remained substantially lower in mutants. (Two-way ANOVA p<.0001 for both metabolites, asterisk denotes p<.05 by Šídák’s multiple comparison test). **(d)** FPLC traces of plasma triglycerides and (**e)** cholesterol were significantly different between wild-type and mutant mice after 20 weeks of exposure to pro-atherosclerotic paradigm. (Two-way ANOVA p<.0001 for both metabolites). (**f)** Brightfield imaging of that aortic arch shows substantial development of atherosclerotic lesions in wild-type, but not mutant, mice, and lesion accumulation was confirmed in (**g)** histological sections and (**h)** *en-face* oil-red-O staining. (**i)** DNA variants near the human *PLA2G12B* gene are significantly associated with plasma triglyceride levels. (**j)** Lead SNPs associated with triglyceride levels are located in regulatory regions in intron 1 of *PLA2G12B*. Red denotes active promoter, yellow denotes weak enhancer, and green denotes genic enhancer in the ChromHMM track.

Major arteries were inspected for signs of atherosclerosis following 20 weeks of exposure to the atherogenic paradigm. Atherosclerotic lesions were readily apparent in the aortic arch and dorsal aorta of wild-type mice, but were virtually undetectable in *Pla2g12b* mutants (Figure 6F). Histological sections (Figure 6G) and *en face* Oil-Red-O staining (Figure 6H) further confirmed the presence of atherosclerotic lesions in wild-type, but not mutant, mice.

Further, we identified two independent genome-wide association studies (Ripatti et al., 2020, Richardson et al., 2020) that associated polymorphisms linked to the *PLA2G12B* promoter with plasma triglyceride levels (Figure 6I). We found that these variants are located in the first intron of *PLA2G12B*, and overlap with a variety of open-chromatin marks and predicted transcription factor binding-sites (Figure 6J). These observations suggest that these human genetic variants may affect triglyceride levels by modulating transcription of *PLA2G12B*.

## DISCUSSION

Over 500 million years ago at the base of the vertebrate lineage, MTP acquired the ability to transfer triglycerides into the ER lumen (Rava and Hussain, 2007, Wilson et al., 2020). While this adaptation laid the foundation for efficient secretion of lipids via the lipoprotein pathway (Figure 7), many additional adaptations were required to produce and secrete mature TRLs. A variety of adaptations enabling secretion of large cargoes such as TRLs have been identified (Ginsberg, 2021, Wang et al., 2021), yet very little is known about how lipid partitioning within the ER lumen is regulated (Shin et al., 2019). Specifically, MTP is capable of transferring lipids to either ApoB-free lumenal lipid droplets or to ApoB-positive nascent TRLs (Wang et al., 2007, Hussain et al., 2003, Sturley and Hussain, 2012), and it is unclear how lipids are apportioned between these two types of particles.

**Figure 7:**
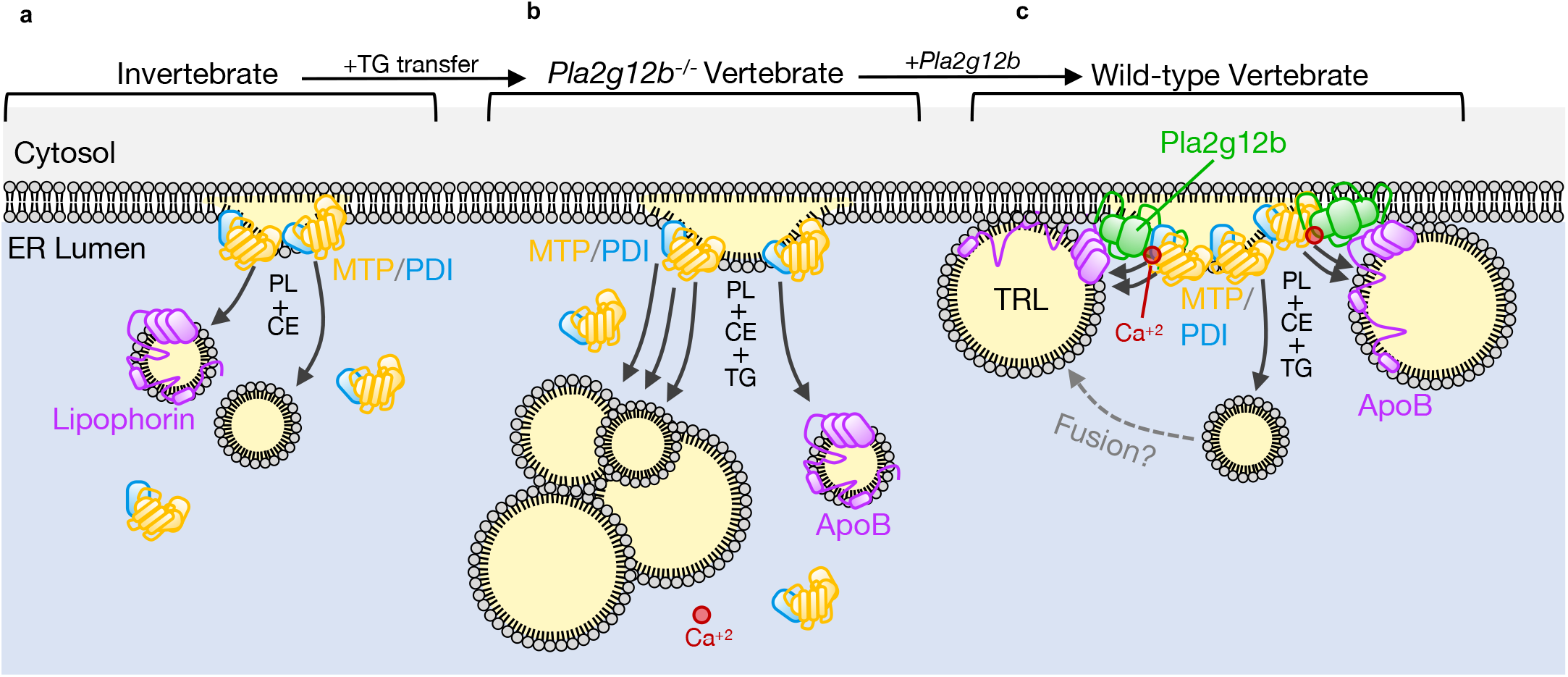
Proposed Model. **(a)** Lipophorin is the invertebrate ortholog of *APOB*, and is capable of forming lipoprotein particles. Invertebrate orthologs of MTP are capable of transferring phospholipids (PL) and Cholesteryl-esters (CE) into the ER-lumen, but are unable to transfer triglycerides (TG). The resulting lipoprotein particles are thus relatively small and dense. (**b)** Vertebrate MTP evolved the capacity to transfer TG into the ER lumen. However, in the absence of functional Pla2g12b, lipids are preferentially transferred to ApoB-free lipid droplets. This results in secretion of lipid-poor lipoproteins and a buildup of unsecretable lipid droplets in the ER-lumen. (**c)** Pla2g12b binds to the lumenal face of the ER-membrane, and also binds calcium, ApoB, and Mtp. These activities promote efficient lipidation of nascent TRLs either through direct transfer or LLD fusion events, and enable production of buoyant TRLs in the vertebrate lineage.

It has also remained unclear why the *pla2g12b* gene has remained conserved in the vertebrate lineage despite having lost its catalytic activity. There are several examples of phospholipase genes losing phospholipase activity and evolving new membranotropic or protein-binding activities (Fujisawa et al., 2008, Ghazaryan et al., 2015), a process called neofunctionalization (Force et al., 1999). Here we address these two longstanding knowledge gaps by identifying Pla2g12b as the first known regulator of lipid partitioning between nascent TRLs and LLDs.

By leveraging the unique advantages of zebrafish, murine, and tissue-culture systems, we were able to gain deep insight into the molecular, cellular, and physiological functions of Pla2g12b. A combination of microscopy and biochemistry techniques allowed us to detect abnormally small lipoproteins and abnormally large lumenal lipid droplets in the ER of *pla2g12b* mutants. We used a panel of chemical probes and secretion assays to show that Pla2g12b is not required for TRL or large-cargo secretion. We then used a novel rescue assay to identify binding partners and annotate functional domains of Pla2g12b, detecting functional interactions with ApoB, Mtp, calcium, and the ER-membrane. Further investigation of the role of calcium in lipoprotein expansion revealed that it is essential for TRL-expansion *in vivo* and *in vitro*, yet dispensable for transfer to ApoB-free lipid droplets. As previously reported, disruption of *PLA2G12B* results in reduction of all classes of serum lipoproteins, including HDL (Aljakna et al., 2012). We suspect this reduction in HDL is secondary to serum hypolipidemia (Chen et al., 2022), but the precise mechanism of this phenotype warrants further investigation. This form of hypolipoproteinemia supported remarkably normal growth and physiology while offering protection from atherosclerosis, suggesting it may have therapeutic or diagnostic value in the prevention of cardiovascular disease.

These observations support a model whereby Pla2g12b acts as a scaffold protein that recruits Mtp and calcium to the lumenal face of the ER-membrane to enhance lipid transfer to nascent TRLs. Both neutral lipids and nascent ApoB-polypeptides are synthesized along the ER-membrane, providing all the necessary building blocks required to assemble a lipoprotein. However, lipids cannot associate with ApoB unless they are transferred by the soluble protein Mtp. Simply recruiting Mtp to the ApoB-rich inner leaflet of the ER-membrane thus represents one mechanism by which Pla2g12b may promote preferential lipid transfer to nascent lipoproteins. We have also shown that hyperphysiological (20mM) calcium supplementation enhances the rate of triglyceride transfer to nascent TRLs, but not LLDs. However, these reactions are performed in *in vitro* conditions that exclude membrane-associated proteins including Pla2g12b. As Pla2g12b contains an essential calcium-binding domain, we hypothesize that Pla2g12b may position calcium ions optimally within the Mtp/ApoB complex to enhance transfer efficiency at physiological calcium concentrations, which approach several hundred micromolar within the ER lumen. Further investigation will be required to determine the precise topology and ion content of this complex, but there is ample evidence to suggest that concentrating ApoB, Mtp, and calcium along the ER-membrane would create a microenvironment that favors lipid transfer to nascent TRLs.

The assays developed here represent a powerful platform for further investigation of TRL biogenesis. We are able to directly visualize subcellular lipid accumulation and monitor expansion of lumenal and serum lipoproteins in parallel, as well as perturb these processes with a panel of genetic and pharmacological probes. The genetic tractability of the YSL offers the opportunity to rapidly interrogate functionality of allelic series using yolk opacity as a simple visual readout. While these assays have greatly advanced our fundamental understanding of sub-organellar lipid trafficking and the poorly characterized gene *pla2g12b*, they also highlight a persistent knowledge gap in our understanding of lipoprotein expansion. Specifically, TRL expansion is thought to occur through both direct transfer of lipids to ApoB by Mtp, and fusion events between TRLs and LLDs (Figure 7C). Although we have proposed the most parsimonious model that supports direct transfer of lipids from the ER to ApoB, we cannot conclusively rule out the existence of an intermediate step whereby Pla2g12b remodels or traffics LLDs to promote fusion with nascent TRLs. Although the net effect on lipoprotein expansion is the same for both models, there is a clear impetus to develop additional tools capable of monitoring LLDs with high spatial and temporal resolution so that we may finally distinguish direct transfer activity from LLD fusion events.

In conclusion, we propose that neofunctionalization of Pla2g12b was a critical event in the evolution of TRLs. Unregulated transfer of lipids into the ER-lumen results in excessive buildup of LLDs, necessitating an additional cofactor to channel lipids to nascent TRLs. We propose that recruitment of ApoB, Mtp, and calcium to sites of neutral lipid synthesis within the inner leaflet of the ER-membrane is sufficient to enhance lipid transfer to TRLs. As a recently duplicated member of a secreted phospholipase gene family, Pla2g12b would have experienced relaxed selective pressure and already possessed membrane and calcium-binding properties, predisposing it to fill this new role. By retaining certain ancestral properties and abandoning others, Pla2g12b emerged at the base of the vertebrate lineage as the master regulator of lipid partitioning between LLDs and TRLs in the ER-lumen.

## ACKNOWLEDGEMENTS

We would like to acknowledge the Mouse Metabolic Phenotyping Core (MMPC) at Vanderbilt University for performing echocardiography, tissue dissections, and FPLC on mouse samples. The Mass Spectrometry Proteomics Core at Baylor College of Medicine provided essential guidance regarding study design and sample preparation for unbiased proteomics and performed mass spectrometry and data analysis. We would also like to thank Sujith Rajan for preparing the unilamellar donor and acceptor particles used in triglyceride transfer assays, Frederick Tan and Allison Pinder for performing RNA-sequencing reactions and data analysis, Kirsten Sadler for providing the ER-TdTomato transgenic line, Michael Matunis for his contribution of superannuated ultracentrifuge supplies that were otherwise unattainable due to supply chain disruptions. This work was financially supported by the NIH (1R01DK093399-09), The Mathers Foundation, and the Carnegie Institution for Science endowment.

## AUTHOR CONTRIBUTIONS

Conceptualization-S.F., J.T. O.F., V.M., M.H., P.Y.; Methodology-J.T., O.F., P.Y.; Investigation-J.T., O.F., P.Y., M.W., T.M., M.S., M.M., K.M.; Resources-E.B.; Writing and Visualization: J.T.; Review and Editing-M.W., O.F., P.Y., T.M., M.S., E.B., M.M., K.M., J.R., V.M., M.H., S.F.; Supervision-J.R., V.M., M.H., S.F.; Funding Acquisition-J.R., V.M., M.H., S.F.

## MATERIALS AND METHODS

### RESOURCE AVAILABILITY

#### Lead contact

Further information and requests for resources and reagents should be directed to and will be fulfilled by the lead contact, Dr. Steven A. Farber.

#### Materials availability

##### Plasmids

Plasmids generated in this study have been deposited to Addgene.

##### Cell Lines

Cell lines generated in this study have been deposited to ATCC.

#### Data and code availability

##### Data

RNA-seq data have been deposited at GEO and are publicly available as of the date of publication. Accession numbers are listed in the key resources table. Microscopy data reported in this paper will be shared by the lead contact upon request.

##### Code

This paper does not report original code.

##### Additional

Any additional information required to reanalyze the data reported in this paper is available from the lead contact upon request.

### EXPERIMENTAL MODEL AND SUBJECT DETAILS

#### Zebrafish

Adult zebrafish were maintained on a 14 h light – 10 h dark cycle and fed once daily with ∼3.5% body weight of Gemma Micro 500 (Skretting USA). All genotypes were bred into the wild-type AB background. As zebrafish sex cannot be determined during the larval stages, sex can be excluded as a variable for experiments involving zebrafish larvae. All procedures comply with all relevant ethical regulations and were approved by the Carnegie Institution Animal Care and Use Committee (Protocol #139).

#### Mice

Adult mice were maintained on a 12 h light – 12 h dark cycle in cages containing ¼” Corn Cob bedding (Envigo, #7097). Mice were provided *ad libitum* access to water and commercially available LM-485 laboratory mouse diet (Envigo, #7012). Mice were housed with littermates whenever possible with densities below 5 adult mice per cage. All procedures comply with all relevant ethical regulations and were approved by the Carnegie Institution Animal Care and Use Committee (Protocol #139).

#### Cells

HepG2 cells were cultured in DMEM 4.5g/L Glucose with UltraGlutamine (Lonza, #H3BE12-604F/U1) medium containing 10% fetal calf serum (Gibco, #10270106), supplemented with 100 units/mL penicillin, and 100 μg/mL streptomycin (Invitrogen, #15070063). Cells were incubated at 37°C in a humidified incubator supplied with 5% CO_2_. Cells were passaged with 0.25% trypsin after reaching 80% confluence. HepG2 cells were seeded in 6-well plates at a density of 5×10^4^ cells/well and grown for two weeks with media replacement every two days. Caco2 cells were grown in the same conditions but in DMEM with 20% FCS, and seeded in Transwell 6-well format dishes (PET, 3.0 µm, Cultek, #45353091), and allowed to differentiate for 21 days under these conditions with changes to fresh medium every 3 days.

### METHOD DETAILS

#### Light Microscopy

Zebrafish larvae were dechorionated manually using forceps prior to imaging when necessary. Larvae were anesthetized in tricaine and mounted laterally in 3% methylcellulose solution (Sigma, M0387) dissolved in embryo medium immediately prior to image collection. Brightfield images were collected using trans-illumination on a Nikon SMZ1500 microscope with HR Plan Apo 1x WD 54 objective set to 3x objective magnification using an Infinity 3 Lumenera camera and Infinity Analyze 6.5 software.

#### Fluorescence Microscopy

Fluorescent images of whole-larvae were collected on a Zeiss Axiozoom v16 microscope set to ×30 magnification equipped with a Zeiss AxioCam MRm.

#### Electron Microscopy

Larvae from each group were fixed, stained with osmium tetroxide, sectioned, and imaged on a Tecnai-12 transmission electron microscope.

#### Microsome preparation

Microsomes were isolated as previously described(Athar et al., 2004) with slight modification. Briefly, 100 larvae were anesthetized in tricaine and homogenized in 1.5 mL chilled buffer A (250 mM sucrose, 50 mM Tris pH 7.4, 5 mM EDTA, .2% Sodium Azide, 1x cOmplete EDTA protease inhibitor cocktail) in a 5 mL Dounce homogenizer. The homogenate was spun at 11k rpm in a chilled benchtop centrifuge to pellet large cellular debris, and the resulting supernatant was adjusted to pH 5.1 with .4 N HCl, rocked at 4C for 10 minutes, and spun at 16k rpm for 30 minutes to pellet microsomes. The resulting pellet was thoroughly resuspended in 1.5 mL buffer B (.054% Sodium Deoxycholate, 1 mM EGTA, 1 mM MgCl_2_, 1 mM Tris pH 7.4) and transferred to a thick-wall polypropylene centrifuge tube (Beckman-Coulter, #349623). Tubes were ultracentrifuged at 50k rpm at 4C in a SW 55 Ti rotor in a Beckman Optima XL-80K Ultracentrifuge. The resulting supernatant was collected as the soluble ER-lumenal fraction.

#### Density Gradient Ultracentrifugation

Density gradient ultracentrifugation was carried out as previously described(Athar et al., 2004) using an Optiprep (D1556, Sigma-Aldrich) gradient optimized to separate lipoprotein subclasses(Yee et al., 2008). Briefly, 1 mL samples were supplemented with 500 uL of Optiprep to yield a 20% iodixanol solution and underlayered beneath 1.5 mL layers of 9% and 12% Optiprep/PBS solutions in a 4.9 mL OptiSeal tube (Beckman-Coulter, 362185) using a disposable glass cannula. Tubes were then ultracentrifuged at 60k rpm for 3 hours in a VTi65.2 rotor in a prechilled Beckman Optima XL 80K Ultracentrifuge at 4C. Optiseal tubes were then punctured and 14 ∼350 uL were collected via drip elution. The refractive index of each fraction was determined using a Bausch and Lomb refractometer, and used to calculate fraction density.

Density profiling of lipoproteins from cultured cells began with induction of VLDL secretion. 1.5ml of Optimem reduced medium without phenol red (Invitrogen: 11058021) supplemented with oleic acid (120nM; Sigma-Aldrich) solubilized in fatty acid–free BSA (Sigma-Aldrich) was added to the cells for 16 hours. After this, media were centrifuged at low speed to remove cells and cellular debris and the resulting supernatant was used directly for the gradients. For cell extracts, cells were washed and collected by scraping in PBS. The cell pellet was resuspended in 1.5ml of hypotonic buffer (10mM Tis-Cl pH 8.8, protease inhibitors), broke with 20 strokes in a cell cracker (EMBL) and incubated for 1 hour on ice. Extracts were centrifuged at 14,000 rpm for 5 min at 4°C. The supernatant was adjusted to 50mm Tris-Cl pH 8.0, 150mM NaCl and used for gradients. 800ul of cleared media or cell extracts were mixed with 200ul of OptiPrep Density Gradient Medium (Stemcell Technologies: 7820). 900ul of the mixture were transferred to a polycarbonate ultracentrifuge tubes (Beckman Coulter:343778), overlayered with 100ul of PBS. Balanced tubes were then loaded into a TLA 120.2 rotor and centrifuged at 115,000rpm in an Optima table-top ultracentrifuge (Beckman Coulter) for 2.5 hours at 16°C. Following ultracentrifugation, density fractions were collected carefully from the top of the tube into 10 separate fractions of 100μL each. The refractive index of each fraction was determined using a refractometer, and used to calculate solution density. 10ul of each fraction, and equivalent amount of media or cell extracts, were immediately mixed with 100ul of PBS and loaded on a Bio-Dot^R^ Microfiltration apparatus (Biorad: 170-6545) for transfer on a nitrocellulose membrane. Dried membranes were blocked with 5% milk and incubated overnight with primary antibodies. After incubation with secondary Alexa-conjugated antibodies, the DotBlot was scanned using Odyssey CLx (LI-COR Bioscience) and individual spots were quantified using QuantityOne (Biorad).

#### Drug treatments

Chemical inhibitors were suspended in DMSO as 2,000X stocks and stored at -20°C: Brefeldin-A (20 mg/mL, Cayman Chemical, #11861), Nocodazole (10 mg/mL, Cayman Chemical, #13857), and Thapsigargin (2 mM, Cayman Chemical, #10522). Working stocks were prepared immediately prior to exposure by supplementing 50 mL of embryo medium with 25 uL of 2,000x stock and 25 uL of DMSO. For dual-drug treatments, the DMSO was replaced with 25 uL of 2,000x stock of the second drug to maintain a constant concentration of 0.1% DMSO. Larvae were incubated in 1x working stock for 90 minutes, and then anesthetized in tricaine and transferred to homogenization buffer for microsome isolation.

#### RNA-Sequencing

Total RNA was isolated from zebrafish larvae using Nucleospin XS micro kit (Macherey-Nagel, #740902.50) following manufacturers instructions. Larvae were collected from three independent clutches, and pooled into groups of 5 for extractions (3 biological replicates per genotype, 5 pooled larvae per replicate). Transcriptome sequencing was then performed using an Illumina NextSeq 500 following library preparation with a Stranded mRNA Library Prep kit (Illumina, #20020594) with TruSeq RNA Single Indexes Set A (Illumina, #20020493).

#### Western Blotting

Media and cell extracts prepared as described above, were mixed with Laemmli SDS sample buffer and denatured for 10 min at 37C, and subjected to SDS-PAGE (6% or 12% acrylamide) followed by Western blotting with the relevant primary antibodies. Following incubation with either Alexa-conjugated or HRP-conjugated secondary antibodies, the blot were imaged using an Odyssey Clx (LI-COR Biosciences) or an Amersham Imager 600 respectively. QuantityOne (Biorad) was used for quantification of individual bands.

#### Triglyceride Transfer Assays

Triglyceride transfer activity was measured *in vitro* using an established method(Athar et al., 2004) scaled down to 384-well plate format. Unilamellar donor vesicles containing nitrobenzoxadiazole (NBD)-labeled TAG as well as un-labeled acceptors were prepared as previously described(Wetterau and Zilversmit, 1984). Each reaction contained 4 uL of donor particles, 4 uL of acceptor particles, 4 uL of mouse microsomal extracts normalized to 400 ug/mL total protein (unless otherwise noted), 4 uL of 10mM Tris-HCl pH 7.4, 4 uL of appropriate salt solution (maximum 200 mM CaCl_2_), and 20 uL of water for a final reaction volume of 40 uL. Microsomal extracts were added last, and after brief mixing by aspiration the plate was transferred to a pre-warmed plate reader set to 37°C and fluorescence was quantified every minute for 2 hours at 460 nm excitation and 530 nm emission. VLDL from human plasma (Lee Biosciences, #365-10) were diluted 50-fold in water and 4 uL of diluted lipoproteins were used in place of standard acceptor vesicles to monitor transfer to lipoprotein particles.

#### Metabolic Profiling

Unbiased metabolic profiling was performed using Dual Energy X-Ray Absorptiometry (DEXA) scans and Comprehensive Lab Animal Monitor Systems (CLAMS). Echocardiograms were performed on conscious mice at baseline and 5 and 10 minute time points following dobutamine injection. Dobutamine was injected IP at a dose of 1.5ug/g body weight. For fractionation of lipoproteins with FPLC, mice were fasted for 4 hours prior to sacrifice, and whole blood was collected in EDTA-coated tubes.

#### Hypercholesterolemia and atherosclerosis induction in mice

Adeno-associated virus constructs (AAV8-D377Y-mPCSK9) were procured from the Vector Biolabs 293 Great Valley Parkway Malvern, PA, USA. These viral vector constructs were injected intravenously (IV) in Wild-type (WT) and *PLA2G12B*^*-/-*^ knock-out mice at the concentration of 10^11^ vector genome copies. Thereafter, mice were fed with a Western diet (WD) for the period of experiment. At the end of the experiment, mice were fasted and mice were euthanized followed by plasma collection and evaluation of atherosclerosis. Plasma was collected to document changes in the plasma triglyceride and cholesterol levels. Size exclusion chromatographic separation of plasma lipoproteins was performed on AKTA pure fast performance liquid chromatography (FPLC). Briefly, 300 μl plasma from WT and KO mice after 6 months of WD feeding were analyzed by FPLC column [Superose™ 6 Increase 10/300 GL FPLC column (GE Healthcare)]. Plasma samples were loaded and then eluted with PBS at a flow rate of 0.4 ml/min and 0.25 ml fractions were collected. Plasma cholesterol and triglyceride levels were measured using a kit (Thermo Fisher Scientific) according to the manufacturer’s protocol. Effect of the Pla2g12b knockout on the cholesterol and triglyceride content in very-low density lipoprotein (VLDL), Intermediate-density lipoprotein (IDL)/low-density lipoprotein (LDL), and high-density lipoprotein (HDL) fractions has been presented in the figure 3. For the evaluation of atherosclerosis, aortas were dissected and examined as per the literature

#### Genotyping

The *pla2g12b*^*sa659*^ allele was genotyped by using primers SF-JHT-400: ACAAGGGAAAGCAAACCAAA and SF-JHT-401: CAGTGTTGTACATGGTGTCTGC to amplify the polymorphic region (57°C T_a_, 30” extension time). The product was subsequently digested with the restriction enzyme BtsI-v2 (NEB, #R0667S), which has a restriction site present only in the sa659 allele.

#### Rescue Injections

Zebrafish embryos were injected at the 1-cell stage with 30 pg of rescue plasmid and 60 pg of capped Tol2 transposase mRNA in an injection mix containing 1 U/uL SUPERase In RNase inhibitor (ThermoFisher Scientific, AM2694) and .05% Phenol Red. At 2 dpf, morphologically normal larvae were screened for GFP expression in the yolk, and GFP-negative larvae were discarded. GFP positive larvae were subsequently anesthetized and imaged as described above and scored for yolk opacity.

#### Co-immunoprecipitation and Mass-spectrometry

Co-immunoprecipitation was performed using the FLAG immunoprecipitation kit (Millipore Sigma, #FLAGIPT1-1KT) following manufacturer instructions. Prior to processing, *pla2g12b*^*-/-*^ larvae were injected with a rescue plasmid containing the *pla2g12b* allele of interest following the rescue injection protocol described above. Injections of ∼300 larvae were generally sufficient to yield at least 50 GFP-positive embryos, which were crosslinked in 2% PFA for 20 minutes at room temperature at 3 dpf. Larvae were homogenized in batches of 10 using pellet pestles in 100 uL of lysis buffer, and the 5 batches were then pooled and the total volume was adjusted to 1 mL. Samples were rocked at 4°C for 30 minutes and ultracentrifuged in a TLA-100 rotor at 44k rpm for 20 minutes to pellet large aggregates. The resulting supernatant was pre-cleared with protein-A coated beads for 2 hours at 4°C, and finally probed with anti-FLAG M2-coated beads overnight at 4C. Beads were submitted to the Mass Spectrometry Proteomics Core at Baylor College of Medicine for protein identification.

#### Generation of PLA2G12B Knock-Out cell lines

To generate Caco2 and HepG2 cell lines in which PLA2G12B expression was abolished, we performed genome editing with the CRISPR/Cas9 system. The online CRISPR Design Tool (https://chopchop.cbu.uib.no/) was used to select two 20-nt guide sequence for pla2g12b gene targeting: sgRNA60 5’GCCAGTACCTACCTGGCTGC3’ and sgRNA94 5’AGGAAGAAGTGATTCCTTCC3’. Construction of the expression plasmid for sgRNA involved a single cloning step with a pair of partially complementary oligonucleotides for each sgRNA. The oligo pairs encoding the 20-nt guide sequences (sgRNA60F 5’CACCGGCCAGTACCTACCTGGCTGC3’ and sgRNA60R 5’AAACGCAGCCAGGTAGGTACTGGCC3’; sgRNA95F 5’CACCGAGGAAGAAGTGATTCCTTCC3’ and sgRNA94R 5’AAACGGAAGGAATCACTTCTTCCTC3’) were annealed and ligated into the plasmid PX458 using the Bbs I cloning site(Ran et al., 2013). The bicistronic pSpCas9 (BB)-2A-GFP (PX458) vector containing cDNAs encoding human codon-optimized Streptococcus pyogenes Cas9 (hSpCas9) with 2A-EGFP, and the remainder of the sgRNA as an invariant scaffold immediately following the oligo cloning site Bbs I, is available from Addgene (plasmid # 48138). Both Caco2 and HepG2 cells were transfected with 1 µg of pooled plasmid using X-tremeGENE9 Transfection Reagent (Roche:6365779001) following manufacturer recommendations. For both cell lines, 96 h after transfection, EGFP-positive cells were isolated by FACS, and single cells were collected in 96-well plates. After expansion to six-well format, differentiated Caco2 and HepG2 cells were collected and protein lysates were prepared to identify lines lacking detectable PLA2G12B protein.

### QUANTIFICATION AND STATISTICAL ANALYSIS

Statistical analyses were performed using Prism software (v9) unless otherwise noted. Details on the statistical tests are described in the associated figure legends. Unless otherwise noted, error bars denote Mean ± Standard Deviation.

## SUPPLEMENTAL INFORMATION TITLES AND LEGENDS

**Supplemental Figure 1: Evolutionary conservation and expression of *pla2g12b*. (a)** Phylogenetic tree of group 12 phospholipase genes demonstrating conservation of both *Pla2g12b* and *Pla2g12a* in every major vertebrate clade. Drosophila sPLA2 is included as an outgroup, as group 12 phospholipases are not detectable outside of the vertebrate lineage. (**b)** Syntenic clusters on human chromosomes 4 and 10 demonstrate that *PLA2G12B* and *PLA2G12A* emerged from a genome duplication event. (**c)** Whole-mount *in-situ* hybridization demonstrating *pla2g12b* expression in the yolk syncytial layer (YSL) in the embryonic stage, and expression in the liver and intestine at later developmental stages. (**d)** qPCR from adult zebrafish intestines showing upregulation of *pla2g12b* gene expression in response to feeding (Welch’s unpaired t-test, asterisk denotes p<.01). (**e)** Schematic representation of sectioning protocol and region of interest for electron micrographs. Transverse sections were cut through the central yolk mass and the syncytium directly adjacent to the yolk mass (YSL) was imaged. (**f)** Brightfield images and corresponding electron micrographs showing an increase in the number of lumenal lipid droplets (dark inclusions) from 2-4 dpf, which correlates with the degree of yolk darkening observed in brightfield images. (**g)** High-magnification electron microscopy suggests that a lipid bilayer membrane surrounds even very large LLDs with diameters >1 µm. (**h)** Epistasis experiment demonstrating that the Mtp inhibitor Lomitapide does impact subcellular lipid accumulation in *pla2g12b* mutants, suggesting that Mtp is upstream of Pla2g12b.

**Supplemental Figure 2: Design of *Pla2g12b* alleles. (a)** Amino acid alignment of Pla2g12b and Pla2g12a homologs from zebrafish, mouse, and human. Amino acids are shaded by similarity, with darker shading indicating higher similarity at that position. (**b)** Legend for labeling scheme in panel a, illustrating that positions that are perfectly conserved (identical) between all proteins are marked with an asterisk. These positions are considered unlikely to be responsible for neofunctionalization of Pla2g12b. Positions that are conserved within orthologs but differ between paralogs are denoted with an orange rectangle. These positions are considered putatively neofunctionalized, as the change is highly conserved in Pla2g12b but absent in Pla2g12a. (**c)** Tabular representation of Pla2g12b alleles. Numbering is based on the position within the zebrafish Pla2g12b amino acid sequence.

**Supplemental Figure 3: Predicted structure of Pla2g12b. (a)** Predicted Structure of Pla2g12b generated by AlphaFold. Ribbon diagram is color-coded based on confidence of the 3D structure, and helix number and direction is indicated by labelled arrows. (**b)** Expected error plot with the essential domains A and B boxed in orange, demonstrating that the spatial relationship between these two domains has very low expected error (dashed orange boxes). (**c)** Space-filling model of Pla2g12b color coded by confidence. (**d)** Space filling model of Pla2g12b with essential domains A and B color coded in Blue and Cyan. (**e)** Space filling model of Pla2g12b color-coded by surface hydrophilicity. (**f)** Structural alignments of Pla2g12b structures from zebrafish, mouse, and human overlap almost perfectly. (**g)** Linear view of predicted domain structure of Pla2g12b, with signal peptide (SP) denoted with a rectangle, helices shown as numbered cylinders, and beta sheets shown as arrows. Domains A and B are denoted with blue and cyan boxes, and the confidence of the predicted structure is shown below (confidence thresholds identical to panel a). Helix numbers and directions correspond to labels in panel a.

**Supplemental Table 1: Differentially expressed genes in *pla2g12b* mutants zebrafish larvae at 4 dpf**. Genes meeting the cutoff threshold for the adjusted p-value and fold change (p<1×10^−5^ and fold-change >1.41) are displayed, sorted by fold change.

**Supplemental Table 2: Unbiased proteomics of Pla2g12b binding partners**. Proteins meeting the cutoff threshold for the adjusted p-value and fold change (p<.15 and fold-change >2) are displayed, sorted by significance. Abundance (Log_2_iBAQ) is also displayed.

**Supplemental Table 3: Protein interactions enriched in functional Pla2g12b isoforms**. The top 50 proteins meeting the cutoff threshold for the adjusted p-value and fold change (p<.05 and fold-change >4) are displayed, sorted by significance. Abundance (Log_2_iBAQ) is also displayed.

