## Supplemental files for "Pla2g12b is Essential for Expansion of Nascent Lipoprotein Particles"

### Supplemental Figure 1: Evolutionary conservation and expression of *pla2g12b*

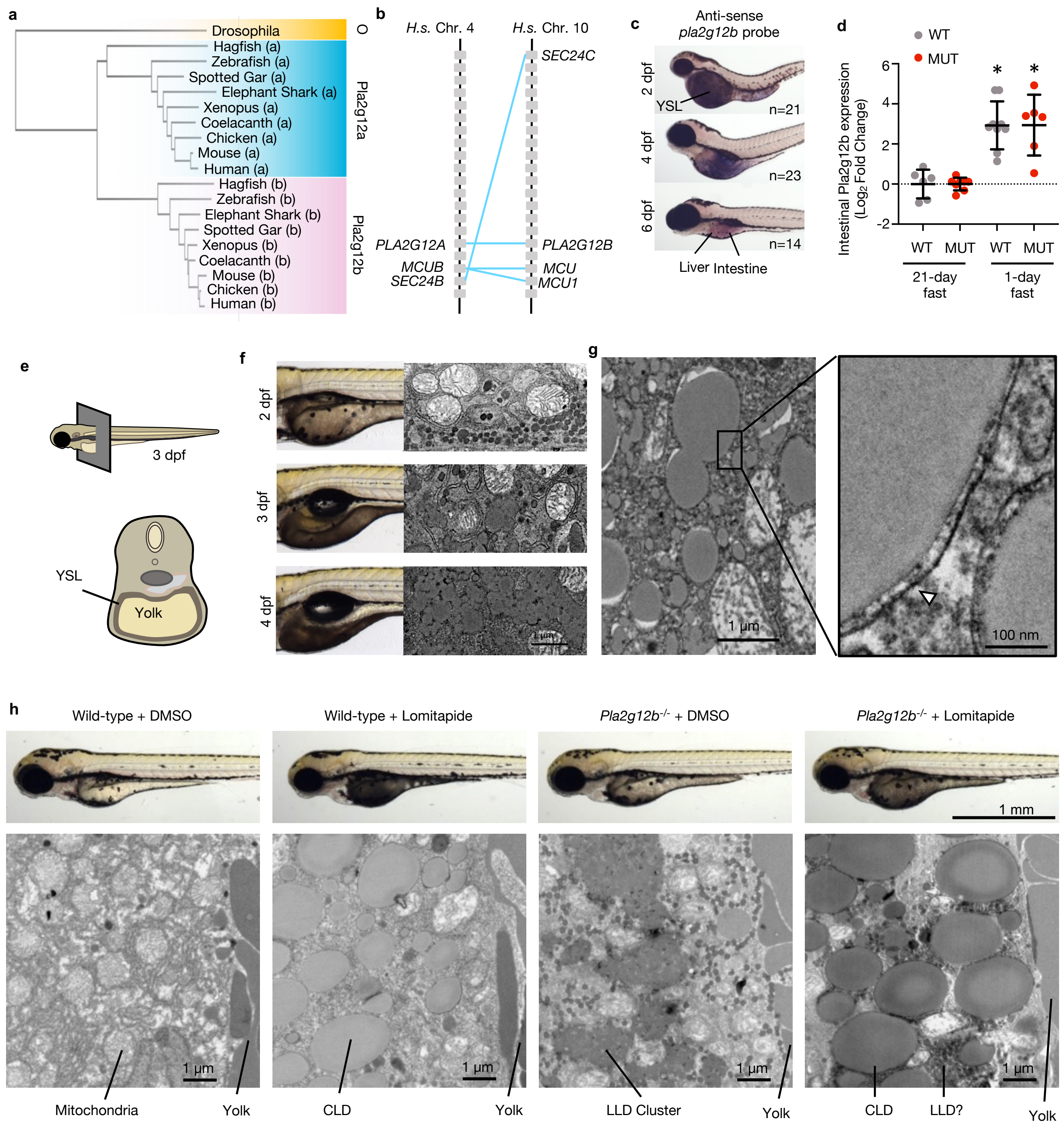

### Supplemental Figure 2: Design of *Pla2g12b* alleles

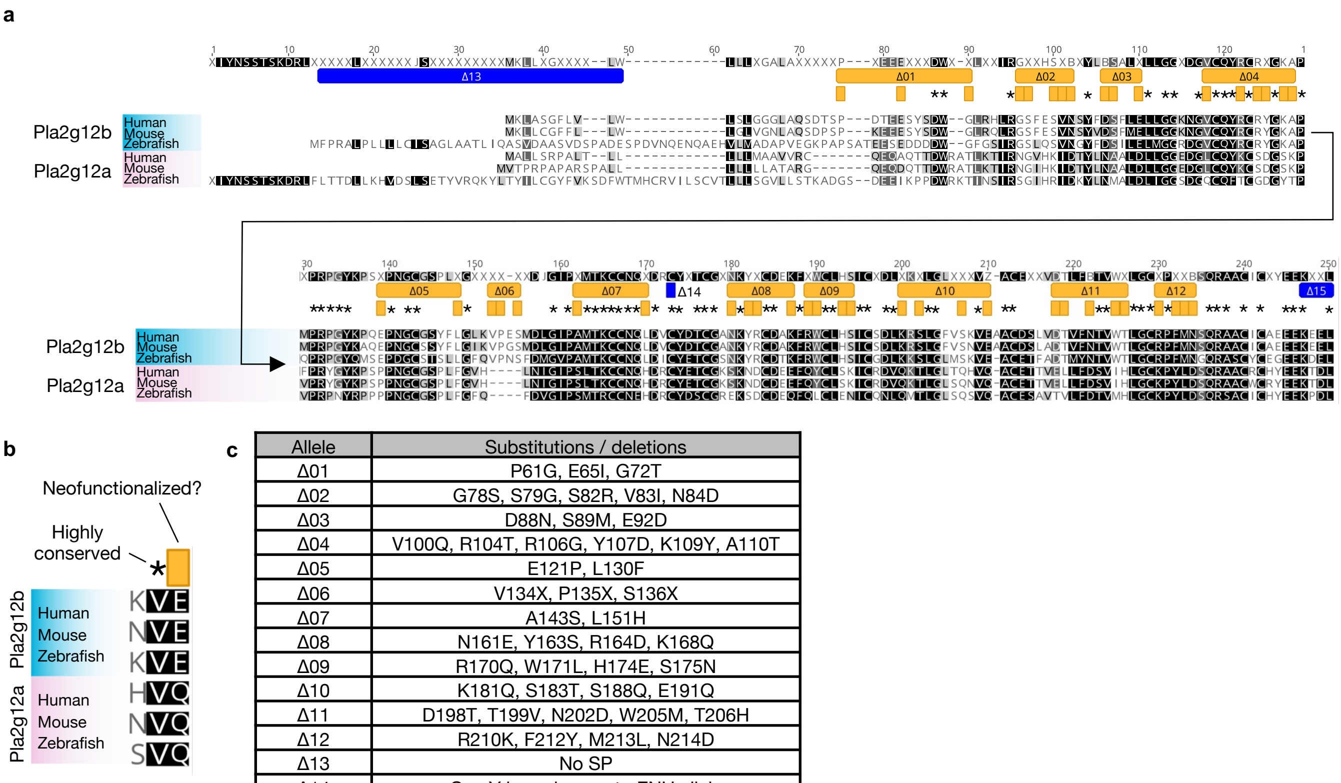


c

| Allele | Substitutions / deletions |
|---|---|
| Δ01 | P61G, E65I, G72T |
| Δ02 | G78S, S79G, S82R, V83I, N84D |
| Δ03 | D88N, S89M, E92D |
| Δ04 | V100Q, R104T, R106G, Y107D, K109Y, A110T |
| Δ05 | E121P, L130F |
| Δ06 | V134X, P135X, S136X |
| Δ07 | A143S, L151H |
| Δ08 | N161E, Y163S, R164D, K168Q |
| Δ09 | R170Q, W171L, H174E, S175N |
| Δ10 | K181Q, S183T, S188Q, E191Q |
| Δ11 | D198T, T199V, N202D, W205M, T206H |
| Δ12 | R210K, F212Y, M213L, N214D |
| Δ13 | No SP |
| Δ14 | C-->Y homologous to ENU allele |
| Δ15 | No KDEL |

### Supplemental Figure 3: Predicted structure of Pla2g12b

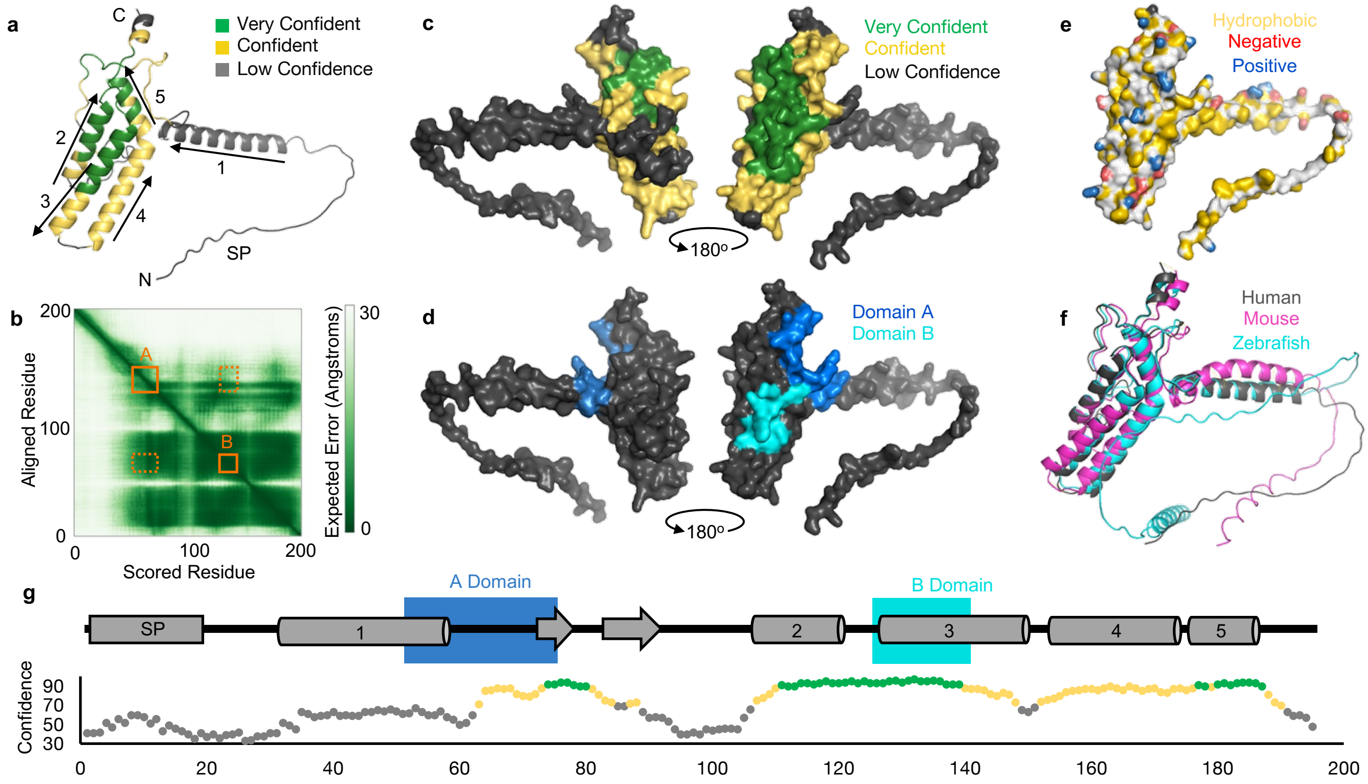

**Supplemental Table 1: Differentially expressed genes in *pla2g12b* mutants zebrafish larvae at 4 dpf**

| names | Geneid | log2FC | padj |
|---|---|---|---|
| cbln11 | ENSDARG00000086654 | 1.891220387 | 7.93E-18 |
| tdo2b | ENSDARG00000023176 | 1.442119458 | 2.92E-07 |
| arg2 | ENSDARG00000039269 | 1.432805551 | 3.82E-19 |
| ucp1 | ENSDARG00000023151 | 1.323555779 | 2.29E-06 |
| c3a.6 | ENSDARG00000043719 | 1.288920897 | 1.23E-08 |
| lpin1 | ENSDARG00000020239 | 1.240153902 | 4.91E-06 |
| hbl4 | ENSDARG00000054202 | 1.192662646 | 6.82E-06 |
| CR855311.1 | ENSDARG00000090352 | 1.189202967 | 1.81E-11 |
| cp | ENSDARG00000010312 | 1.166023778 | 3.79E-06 |
| uroc1 | ENSDARG00000070394 | 1.055899704 | 3.93E-11 |
| mylipa | ENSDARG00000008859 | 1.020959613 | 1.14E-07 |
| abca1a | ENSDARG00000074635 | 1.001697465 | 3.08E-16 |
| c3a.1 | ENSDARG00000012694 | 0.984013829 | 3.69E-09 |
| lpl | ENSDARG00000087697 | 0.974893207 | 2.67E-08 |
| si:ch211-264e16.1 | ENSDARG00000068947 | 0.945473782 | 6.47E-06 |
| pnpla2 | ENSDARG00000089390 | 0.851177682 | 6.47E-06 |
| slc13a2 | ENSDARG00000053853 | 0.803022785 | 4.29E-08 |
| slc25a48 | ENSDARG00000021250 | 0.79732337 | 2.21E-09 |
| psme4a | ENSDARG00000087911 | 0.790868737 | 6.82E-06 |
| hpxb | ENSDARG00000051912 | 0.787322577 | 6.82E-06 |
| nfe2l1a | ENSDARG00000030616 | 0.74297671 | 2.29E-06 |
| ckba | ENSDARG00000069752 | 0.717663246 | 3.42E-07 |
| pah | ENSDARG00000020143 | 0.537824856 | 2.07E-08 |
| itgb1b.2 | ENSDARG00000022689 | 0.522042746 | 9.99E-06 |
| fgfr3 | ENSDARG00000004782 | -0.66957842 | 2.82E-07 |
| zgc:163030 | ENSDARG00000069192 | -0.98245064 | 1.51E-09 |
| pla2g12b | ENSDARG00000015662 | -1.30535546 | 1.41E-14 |
| adam8a | ENSDARG00000001452 | -1.52877021 | 1.81E-11 |
| cyp2aa8 | ENSDARG00000104540 | -1.53460217 | 3.18E-06 |
| sqlea | ENSDARG00000079946 | -1.58198026 | 2.35E-08 |
| gck | ENSDARG00000068006 | -2.78426898 | 1.53E-12 |

**Supplemental Table 2: Unbiased proteomics of Pla2g12b binding partners**

| Gene Symbol | t-test | log2-fc | log2-abundance |
|---|---|---|---|
| dnajc3a | 1.14976E-06 | 20.68243271 | 7.39472033 |
| pla2g12b | 7.65426E-05 | 20.65896745 | 22.26805782 |
| apobb.1 | 0.030499149 | 3.399952677 | 17.77433903 |
| p4hb | 0.044879762 | 4.284491852 | 18.55466534 |
| arcn1b | 0.047649592 | 13.90972832 | 13.52799429 |
| hspa5 | 0.070280042 | 2.332962656 | 19.73098657 |
| nt5c2b | 0.083238051 | 5.044439905 | 13.12123671 |
| prdx4 | 0.086358992 | 4.918892077 | 15.41631984 |
| arf4a | 0.086790542 | 5.170837761 | 11.91208533 |
| hsp90b1 | 0.088556229 | 3.266685512 | 17.8331628 |
| hyou1 | 0.096853589 | 4.532544039 | 14.4857246 |
| ube2d3 | 0.104045528 | 2.565240254 | 11.71611056 |
| ube2d2l | 0.104045528 | 2.565240254 | 11.71611056 |
| ube2d2 | 0.104045528 | 2.565240254 | 11.71611056 |
| rac1l | 0.106494473 | 3.395118259 | 10.30299884 |
| tpi1a | 0.116116524 | 14.84076554 | 1.553053156 |
| magt1 | 0.116118935 | 22.74993639 | 9.462224007 |
| trappc12 | 0.116119797 | 19.84150742 | 6.553795036 |
| atf1 | 0.116120058 | 19.37231354 | 6.08460116 |
| crema | 0.116120058 | 19.37231354 | 6.08460116 |
| creb1b | 0.11612012 | 19.20519092 | 5.917478539 |
| tecrb | 0.116127403 | 19.10360528 | 5.815892901 |
| macrod1 | 0.116141882 | 17.66764519 | 4.379932812 |
| LOC100537744 | 0.116145026 | 26.09451107 | 12.80679869 |
| hsd17b12b | 0.116147253 | 23.46483523 | 10.17712285 |
| adssl | 0.116147632 | 20.94026063 | 7.652548246 |
| ttc7b | 0.116154256 | 21.91623361 | 8.62852123 |
| appl1 | 0.116240585 | 22.31687954 | 9.029167164 |
| uggt1 | 0.116242012 | 24.10428774 | 10.81657536 |
| vapal | 0.116270527 | 24.3904316 | 11.10271923 |
| slc25a6 | 0.116285288 | 25.99153982 | 12.70382744 |
| gde1 | 0.116290107 | 18.44923433 | 5.161521954 |
| tsta3 | 0.116308941 | 17.83347999 | 4.545767607 |
| mrps17 | 0.116352273 | 12.60869491 | 13.61200399 |
| mccc2 | 0.116404117 | 24.87578952 | 11.58807714 |
| rsu1 | 0.116526578 | 22.13597713 | 8.848264751 |
| pitpnb | 0.116568154 | 20.14291055 | 6.855198175 |
| adssl1 | 0.116620594 | 24.82319979 | 11.53548741 |
| tuba1a | 0.116647715 | 34.56355431 | 21.27584193 |
| nudt9 | 0.116965206 | 24.82264761 | 11.53493523 |
| eif4a1b | 0.117173289 | 29.53318885 | 16.24547647 |
| calb2b | 0.117317102 | 25.90344313 | 12.61573075 |
| h6pd | 0.117512259 | 20.05276685 | 6.765054471 |
| hdac10 | 0.117999452 | 19.17405472 | 5.886342341 |
| zgc:136908 | 0.118170595 | 23.90478103 | 10.61706865 |
| zgc:112271 | 0.120454202 | 1.338251142 | 14.10519099 |
| hspa4a | 0.121514621 | 22.98980329 | 9.702090912 |
| rps26l | 0.125324073 | 1.354704349 | 15.62628349 |
| mttp | 0.134528211 | 8.328166776 | 17.9536142 |

**Supplemental Table 3: Protein interactions enriched in functional Pla2g12b isoforms**

| Gene Symbol | t-test | log2-fc | log2 abundance |
|---|---|---|---|
| pdxka | 0.000170716 | 13.72648454 | -3.75 |
| lsm3 | 0.000170716 | 13.70534007 | -3.76 |
| smx5 | 0.000170716 | 13.77382306 | -3.72 |
| ugt1b7 | 0.000258955 | 2.072996702 | 0.68 |
| hddc3 | 0.000403471 | 3.794108929 | -7.22 |
| enkur | 0.000528479 | 4.758395953 | 2.55 |
| ppp1r3b | 0.000544723 | 4.567738349 | 2.40 |
| rab3gap1 | 0.000577513 | 4.272447314 | 2.11 |
| snrpg | 0.000592268 | 1.978404272 | 2.02 |
| ssrp1a | 0.000614452 | 4.01125004 | 1.91 |
| usp4 | 0.000617635 | 4.180098424 | 1.95 |
| mgea5l | 0.000685619 | 3.533232151 | 1.50 |
| vim | 0.000848288 | 9.93364739 | 2.67 |
| pbrm1 | 0.001473794 | 4.260577648 | 3.56 |
| nrbp1 | 0.001822778 | 1.927150269 | 1.12 |
| eif3jb | 0.001864544 | 1.993061619 | 1.15 |
| mgll | 0.001897982 | 2.043053344 | 1.15 |
| hyi | 0.001921272 | 2.077666704 | 1.29 |
| sars | 0.001948433 | 2.120882666 | 1.13 |
| si:dkey-11k2.7 | 0.002047244 | 5.490853516 | -1.34 |
| nubp2 | 0.002054503 | 2.284456485 | 1.19 |
| tomm40 | 0.002162742 | 2.450655207 | 1.19 |
| endou2 | 0.002505937 | 3.420622064 | 1.08 |
| ralaa | 0.002676458 | 3.336902772 | 1.15 |
| ralbb | 0.002676458 | 3.336902772 | 1.15 |
| ralba | 0.002676458 | 3.336902772 | 1.15 |
| ralab | 0.002676458 | 3.336902772 | 1.15 |
| atox1 | 0.002706558 | 3.717270772 | 1.11 |
| banf1 | 0.002920883 | 4.053320849 | 1.20 |
| ctbp2a | 0.003290335 | 1.552411808 | -3.73 |
| zgc:136908 | 0.003519295 | 2.676574497 | -13.29 |
| si:ch1073-159d7.7 | 0.003567813 | 4.633355518 | -13.29 |
| si:dkey-23a13.17 | 0.003567813 | 4.633355518 | -13.29 |
| si:dkey-23a13.9 | 0.003567813 | 4.633355518 | -13.29 |
| zgc:110425 | 0.00361504 | 5.464366097 | -13.29 |
| blmh | 0.003898083 | 2.801431412 | 0.87 |
| isoc1 | 0.004044133 | 4.945708164 | 1.09 |
| hrasb | 0.004597007 | 9.688509709 | 1.82 |
| LOC100538266 | 0.00598882 | 4.01489789 | -5.58 |
| nme2a | 0.006090459 | 4.10551881 | -5.56 |
| zgc:162816 | 0.006322383 | 4.365820907 | 0.41 |
| capn8 | 0.006385232 | 4.453964103 | 0.43 |
| nmnat1 | 0.00646669 | 4.199288642 | 0.29 |
| mapkapk2a | 0.006472676 | 4.219072106 | 0.34 |
| nat16 | 0.006755941 | 3.878839509 | 0.05 |
| atp2b4 | 0.006828565 | 3.797426101 | -0.01 |
| ctsz | 0.007258097 | 3.387096311 | -0.31 |
| ogdhb | 0.008311155 | 19.06707959 | -4.34 |
| qkib | 0.008475538 | 4.116605612 | 4.43 |
